# GRK-dependent ACKR3 endocytosis and chemokine scavenging is independent of receptor phosphorylation and β-arrestin

**DOI:** 10.64898/2026.05.11.724365

**Authors:** Boubacar Sidiki Traore, Sabrina Casella, Pierre Couvineau, Meriem Semache, Diego Morone, Gianluca D’Agostino, Sylvia Thelen, Billy Breton, Pedro Henrique Scarpelli Pereira, Mariagrazia Uguccioni, Daniel F. Legler, Marcus Thelen, Michel Bouvier

## Abstract

Desensitization and internalization of most G protein-coupled receptors (GPCRs) depend on phosphorylation by GPCR kinases (GRKs), promoting β-arrestin recruitment. Atypical chemokine receptors (ACKRs), including ACKR3, are structurally related to classical chemokine receptors but do not activate heterotrimeric G proteins. ACKR3 signaling and trafficking have been proposed to depend on GRK5-mediated phosphorylation and β-arrestin interaction. However, the respective roles of β-arrestins, GRKs, and receptor phosphorylation in chemokine scavenging and in constitutive or ligand-induced trafficking remain debated. Using bioluminescence resonance energy transfer (BRET)–based biosensors and immunofluorescence imaging with fluorescently labeled receptors and chemokines, we examined ACKR3 interaction with β-arrestin1/2 and assessed chemokine scavenging and receptor trafficking in β-arrestin–deficient (Δβarr1/2) cells. We also evaluated the contribution of GRK-mediated phosphorylation. β-arrestins supported agonist-independent receptor internalization but were dispensable for chemokine-induced internalization and chemokine scavenging. In contrast, GRKs were required for ligand-promoted endocytosis, with either GRK2/3 or GRK5/6 being sufficient. Mutation of ACKR3 phosphorylation sites impaired β-arrestin recruitment but did not completely block internalization and scavenging, whereas complete C-terminal truncation abolished both processes. Consistently, kinase-dead GRK2 rescued ACKR3 endocytosis in ΔGRK2/3/5/6 cells, indicating a scaffolding role partially independent of kinase activity. Moreover, Gβγ was not required for GRK2-mediated ACKR3 endocytosis, as a PH-domain–deleted GRK2 mutant restored internalization in ΔGRK2/3/5/6 cells, and Gβγ sequestration by βARKct-CAAX did not inhibit this process consistent with the notion that ACKR3 does not promote G protein activation. Thus, ligand-promoted ACKR3 internalization and chemokine scavenging occur independently of β-arrestins but requires GRKs.

**One-sentence summary:** GRKs are essential for ACKR3 endocytosis and chemokine scavenging, whereas β-arrestins and receptor phosphorylation are dispensable.

## Introduction

Chemokines together with their cognate receptors constitute the chemokine system that is critically involved in leukocyte migration in homeostatic and inflammatory conditions [1]. In addition, the chemokine system can guide cell trafficking during development, tissue remodeling, tumorigenesis and metastasis [2–4]. Cell migration is triggered through complex signaling cascades induced by binding of a chemokine to its typical chemokine receptors. Typical or classical chemokine receptors belong to the γ-subclass of rhodopsin-like G protein-coupled receptors (GPCRs) [5]. Directional movement requires the presence of a chemotactic gradient on which a cell can orient itself. Gradients are formed by the secretion of chemokines, often by endothelium and stromal cells, and their retention by glycosaminoglycans (GAGs) present on the extracellular matrix or cell surfaces. In addition, atypical chemokine receptors (ACKRs) play a pivotal role in carving and maintaining the gradients [6–8]. Because ACKRs do not couple to G proteins, chemokine binding does not lead to the activation of heterotrimeric G proteins and hence, does not trigger cell migration [9, 10]. ACKRs mainly function as scavengers cycling spontaneously and in response to chemokines, between the plasma membrane and endocytic compartments where they deliver their cargo for lysosomal degradation before cycling back to the plasma membrane [11]. The ACKR-mediated ligand-depletion controls chemokine availability in the extracellular milieu, contributing to local gradient formation, which in turn favors cell migration [12]. ACKR3 (also known as CXCR7) is a scavenging receptor for the chemokines CXCL12 and CXCL11, which bind and activate the canonical receptors CXCR4 and CXCR3, respectively [13–15]. In line with the scavenger function, several studies show that ACKR3 activity supports the formation of CXCL12-gradients in embryonic development [16–19].

ACKR3 cycles spontaneously between the plasma membrane and endosomal compartments, but the process can be enhanced by ligands [15]. Genetic ablation of ACKR3 in C57BL/6 mice, which do not express CXCL11, leads to perinatal death [20], suggesting that ACKR3-mediated scavenging of CXCL12 is critical for survival. However, it was also reported that a knock-in variant of ACKR3 lacking all the receptor’s C-terminal phosphorylation sites, which shows a markedly reduced ability to scavenge CXCL12 is fully viable [21].

β-arrestin1 and 2 (also called arrestin-2 and arrestin-3, respectively) regulate the signaling of many GPCRs mediating receptor desensitization and internalization and can promote signaling activities on its own [22, 23]. The recruitment of these cytosolic proteins is triggered by agonist-induced phosphorylation of serine/threonine residues at the C-terminus, the third intracellular loop or of both domains of the receptors by second messenger kinases and G protein-coupled receptor kinases (GRKs) [24–26]. ACKRs, except for ACKR1, have been reported to recruit β-arrestin1 and 2 and may trigger β-arrestin-dependent signaling [27–30]. ACKR2, ACKR3, ACKR4 and ACKR5 have been proposed to be constitutively associating with β-arrestins, which could account for spontaneous receptor trafficking [15, 31–35]. Moreover, the scavenging activity of ACKRs was attributed to β-arrestin recruitment promoting receptor endocytosis [28, 31, 36, 37]. Several studies have challenged the role of interaction between ACKR3 and β-arrestins, questioning the necessity of this adaptor protein in CXCL12-mediated internalization. The main observation was that ACKR3-mediated chemokine scavenging still occurs in the absence of β-arrestins [30, 38]. Moreover, Saaber *et al*. suggested that, although requiring GRK2-mediated phosphorylation of its C-terminus, ACKR3-mediated chemokine depletion is β-arrestin independent [21]. An exclusive role of GRK5/6 and not GRK2/3 has been suggested to be important for the recruitment of β-arrestin1 and-2 [39]. In other studies, a predominant role for GRK5 rather than GRK2 has been proposed to be responsible for receptor internalization and chemokine scavenging; only the GRK5-mediated phosphorylation resulting directly from ACKR3 activation whereas GRK2-mediated phosphorylation was suggested to result from cross-activation of CXCR4 by CXCL12 [40]. More recently, *in vitro* phosphorylation assays coupled to cryo-EM structures revealed that phosphorylation by each kinase resulted in different receptor-β-arrestin complexes, suggesting potential different outcomes form the β-arrestin interaction promoted by each kinase [41]. It follows that the relative roles of receptor phosphorylation, β-arrestins and specific GRKs in the constitutive and ligand promoted receptor endocytosis and chemokine scavenging remain debated and deserve further characterization.

Here, using a combination of imaging, bioluminescence resonance energy transfer and genetic engineering approaches, we systematically assessed β-arrestin1 and 2 recruitments to ACKR3 as well as their roles in the constitutive and ligand-promoted receptor trafficking as well as chemokine scavenging. Our results show that these adaptor proteins support constitutive receptor internalization but confirm the proposition that they are dispensable for ligand-promoted trafficking and chemokine scavenging. We found that such endocytosis requires either GRK2/3 or GRK5/6 family members but that this action of the kinases is partially phosphorylation independent.

## Results

### ACKR3 trafficking in the absence of β-arrestins

Many GPCRs require β-arrestins for internalization [42]. Given that ACKR3 precouples to β-arrestin2 [43] and can recruit β-arrestins upon chemokine binding, and given the contradictory reports concerning their roles in ACKR3 endocytosis [30, 36, 44, 45], we asked whether ACKR3 can be internalized in the absence of β-arrestins. To this end we used HeLa cells in which both β-arrestins (Δβarr1/2) were deleted by gene editing [46] and measured spontaneous as well as chemokine-dependent internalization of ACKR3. To measure accurately ACKR3 trafficking we used the chimeric chemokine CXCL11-12 to avoid interference of CXCL12 binding to CXCR4 that is expressed at low levels in most cell types, including HeLa cells [47], and that could explain some of the differences observed between studies. The chimera is composed of the N-terminus of CXCL11 and the main body of CXCL12, and was shown to selectively bind to ACKR3 with high affinity, but - unlike CXCL12 and CXCL11 - does not bind to the chemokine receptors CXCR4 and CXCR3 [48]. HeLa cells were transiently transfected with an ACKR3 construct that was tagged at the N-terminus with the S6 peptidyl carrier protein consensus sequence (PCP-ACKR3), which has high specific activity for the Sfp phosphopantetheinyl transferase (PPTase) [49, 50]. Cell surface expressed PCP-ACKR3 was labeled with Coenzyme A conjugated with Atto647N at 17°C, to reduce receptor internalization during labeling [50]. We determined the internalization of Atto647-PCP-ACKR3 in wild-type and Δβarr1/2 HeLa cells (Figure 1A) in the presence and absence of CXCL11-12 using confocal microscopy. In agreement with previous findings [50], imaging of cells fixed immediately after labeling at 17°C showed a clear plasma membrane localization of surface labelled PCP-ACKR3 in wild-type and Δβarr1/2 cells (Figure 1A, upper panels). No obvious differences of ACKR3 compartmentalization were observed at this time point whether β-arrestins were present or not. Shifting the cells to 37°C for 8 min caused the spontaneous internalization of surface labelled PCP-ACKR3 in the absence of ligand for both wild-type and Δβarr1/2 HeLa cells (Figure 1A, middle panels), consistent with a robust ligand-independent receptor endocytosis that does not require β-arrestins. Similarly, in the presence of CXCL11-12 surface labelled PCP-ACKR3 readily internalized in wild-type and Δβarr1/2 HeLa cells (Figure 1A, lower panels). To assess more quantitatively possible contributions of β-arrestins, we measured the levels of ACKR3 present at the plasma membrane in wild-type and Δβarr1/2 HeLa cells following spontaneous or ligand-promoted internalization. For this purpose, plasma membranes were stained with Vybrant™ DiO to define the perimeter of single cells. The outer perimeter was drawn manually, consecutive 1 µm large regions of interest (ROIs) were calculated automatically and analyzed for ACKR3 associated fluorescence using ImageJ (Supplementary Figure 1). Cumulative analysis of 25-36 cells revealed in wild-type cells a constitutive endocytosis of ACKR3. A small, but significant difference of spontaneous ligand-independent internalization of ACKR3 was observed between wild-type and Δβarr1/2 HeLa cells (Figure 1B). In the absence of β-arrestins, spontaneous receptor internalization was reduced, resulting in 35% more receptor at the cell surface in the Δβarr1/2 cells following 8 minutes at 37C. The data indicate that both, spontaneous ACKR3 internalization and chemokine scavenging occur efficiently in the absence of β-arrestins, but suggest that the presence of β-arrestins may contributes to spontaneous internalization.

**Figure 1.**
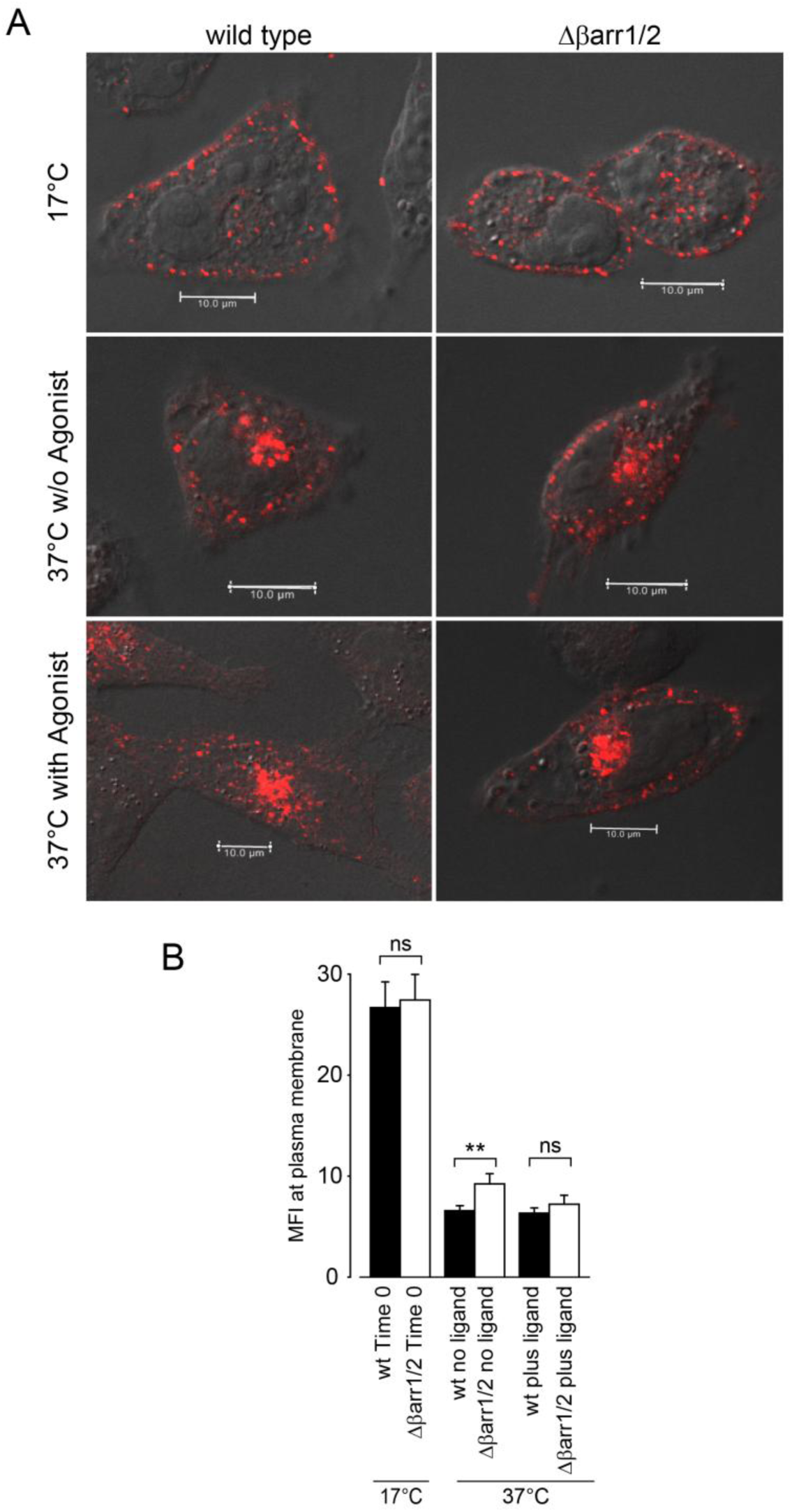
ACKR3 internalization in the presence and absence of β-arrestins. **(A)** Wild-type (left panels) and Δβarr1/2 (right panels) HeLa cells were transfected with a plasmid encoding PCP-ACKR3 and labeled at 17°C with CoA-Atto-647N (red). Shown are the confocal images of PCP-ACKR3 (red) immediately after labeling at 17°C (upper panels) and after 8 min at 37°C (middle panel). For the cells shown in the lower panels, 50 nM CXCL11-12 was added for 8 min at 37°C. Gray: phase contrast. Scale bar 10 µm. (B) Cells expressing PCP-ACKR3 were labeled at 17°C and fixed immediately (Time 0) or incubated at 37°C for 8 min in the absence (no ligand) or presence of 50 nM CXCL11-12 (plus ligand). Mean fluorescence intensity at the plasma membrane was measured with a reduced pinhole (0.8 Airy) to confine signals to the outer perimeter of the cells. Means ± SEM from 25-36 cells. Statistical analysis was performed with ordinary one-way ANOVA corrected with Tukey’s multiple comparisons test (ns: not significant, **p<0.012).

### Subcellular localization of ACKR3 and CXCL11-12

To monitor chemokine scavenging, the subcellular localization of Atto565-labeled CXCL11-12 (CXCL11-12AT565) was assessed simultaneously with Atto647-PCP-labeled ACKR3 in wild-type and Δβarr1/2 HeLa cells. Figure 2A shows a marked spontaneous receptor internalization (upper panels) and chemokine uptake (lower panels) of fluorescently labeled PCP-ACKR3 in wild-type (left) and Δβarr1/2 HeLa (right) cells. In the presence of Atto565-labeled CXCL11-12 (CXCL11-12AT565) [48], the receptor and the chemokine similarly colocalized in endosomal structures in both cell backgrounds. To further characterize the localization of ACKR3, rGFP constructs fused to the polybasic sequence and prenylation CAAX box of K-Ras (rGFP-CAAX) or the FYVE domain of endofin (rGFP-FYVE) were used as markers of the plasma membrane and early endosome respectively [51]. As shown in Figure 2B, when fluorescently labeled PCP-ACKR3 expressed in wild-type or Δβarr1/2 HEK293 cells were shifted to 37°C for 10 min in the presence of CXCL11-12AT565 [48], most of the receptor originally labelled with Atto647N at the cell surface colocalized with the rGFP-FYVE (right panels) and not the rGFP-CAAX (left panels). Receptor-ligand complexes were exclusively located in endosomes harboring rGFP-FYVE (right panels) in both cell backgrounds, indicating that β-arrestins are dispensable for ACKR3-mediated chemokine uptake.

**Figure 2.**
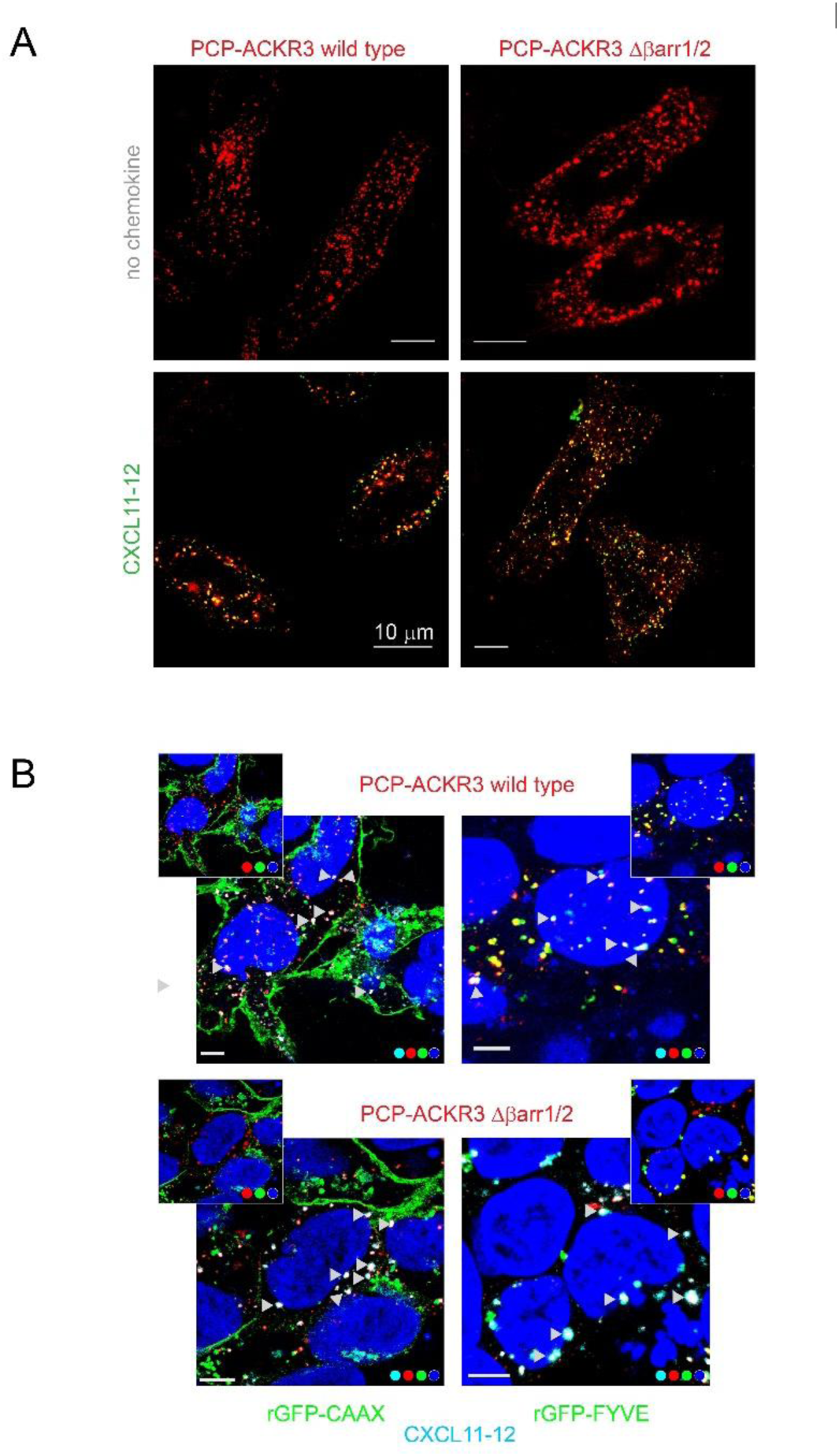
ACKR3 mediated chemokine scavenging in the presence and absence of β-arrestins. **(A)** Wild-type (left panels) and β-arrestin depleted (Δβarr1/2, right panels) HeLa cells were transfected with a plasmid encoding PCP-ACKR3 and enzymatically labeled at 17°C with CoA-Atto647N (red) for 20 min. Cells were incubated for 10 min at 37°C in the absence of chemokine (upper panels) or in the presence of 50 nM fluorescently labeled CXCL11-12Atto565 (lower panels, green). Confocal images show colocalization of fluorescent labeled ACKR3 (red) and CXCL11-12 (green) in yellow. Scale bar in the images 10 µm. (B) Wild-type (upper panels) and β-arrestin depleted (Δβarr1/2, lower panels) HEK293 cells were co-transfected with plasmids encoding rGFP-CAAX (green, left panels) or rGFP-FYVE (green, right panels) as well as PCP-ACKR3 enzymatically labeled at 17°C with CoA-Atto647N (red) and incubated for 10 min at 37°C in the presence of 50 nM CXCL11-12Atto565 (cyan). Receptor-ligand complexes (white: red and cyan) are marked with arrow heads. Images were taken by confocal microscopy. Nuclei were stained with DAPI (blue). For clarity in the smaller frames (upper corners) the cyan channel was turned off showing colocalization of ACKR3 and GFP in yellow, in the large panels CXCL11-12Atto565 is visible. Scale bar in the large images 5 µm.

### Ligand-dependent trafficking of ACKR3

Given that conflicting data exist on the role of β-arrestins in ACKR3 trafficking [30, 36, 44, 45], the ligand-promoted endocytosis of ACKR3 was further monitored using quantitative bioluminescence resonance energy transfer assays. For this purpose, WT and Δβarr1/2 HEK293 cells were transiently transfected with ACKR3-RlucII as BRET-donor and with rGFP-CAAX or rGFP-FYVE as acceptors. CXCL12 induced a time-dependent translocation of ACKR3 from the plasma membrane (PM) to endosomes as illustrated by the decrease in BRET between ACKR3-RlucII and rGFP-CAAX (Figure 3; left panels, CAAX) and a concomitant increase in BRET between ACKR3-RlucII and rGFP-FYVE (Figure 3; right panels, FYVE). The kinetic of ACKR3 translocation to endosomes was very similar between wild-type (WT) and Δβarr1/2 HEK293 cells and the re-expression of either β-arrestin1 or β-arrestin2 did not affect the endocytosis profiles (Table 1). Interestingly, the absence of β-arrestins resulted in a slower disappearance from the CAAX-positive PM domain that was corrected following the reintroduction of either β-arrestin1 and β-arrestin2 (Table 1), suggesting that whereas β-arrestin is dispensable for ACKR3 endocytosis, it may contribute to its trafficking kinetics. As controls we used the chemokine receptor CXCR4 and the mu-opioid (MOR) receptors that have previously been reported to undergo endocytosis in a β-arrestin-independent [52, 53] and -dependent [53] manner, respectively. As expected and seen in Supplementary Figure 2A, the CXCL12-promoted endosomes translocation profile of CXCR4 was similar in WT and in Δβarr1/2 HEK293 cells but in contrast with ACKR3, the re-expression of either β-arrestin1 or β-arrestin2 increase the extent of endocytosis indicating that for this receptor β-arrestins can contribute to some extent to the steady-state endocytosis. In Supplementary Figure 2B, the met-enkephalin-induced MOR endocytosis was completely abolished in the Δβarr1/2 cells. Reintroducing either β-arrestin1 or β-arrestin2 restored an endocytic kinetic almost identical to that observed in the WT cells. A similar β-arrestin-independent ACKR3 endocytosis was observed in HeLa cells in response to either CXCL12 or CXCL11-12 (Supplementary Figure 3A). Taken together these data confirm that ACKR3 can undergo robust endocytosis in the absence of β-arrestins in different cell types and that β-arrestins may have a role in the kinetic of this process.

**Figure 3.**
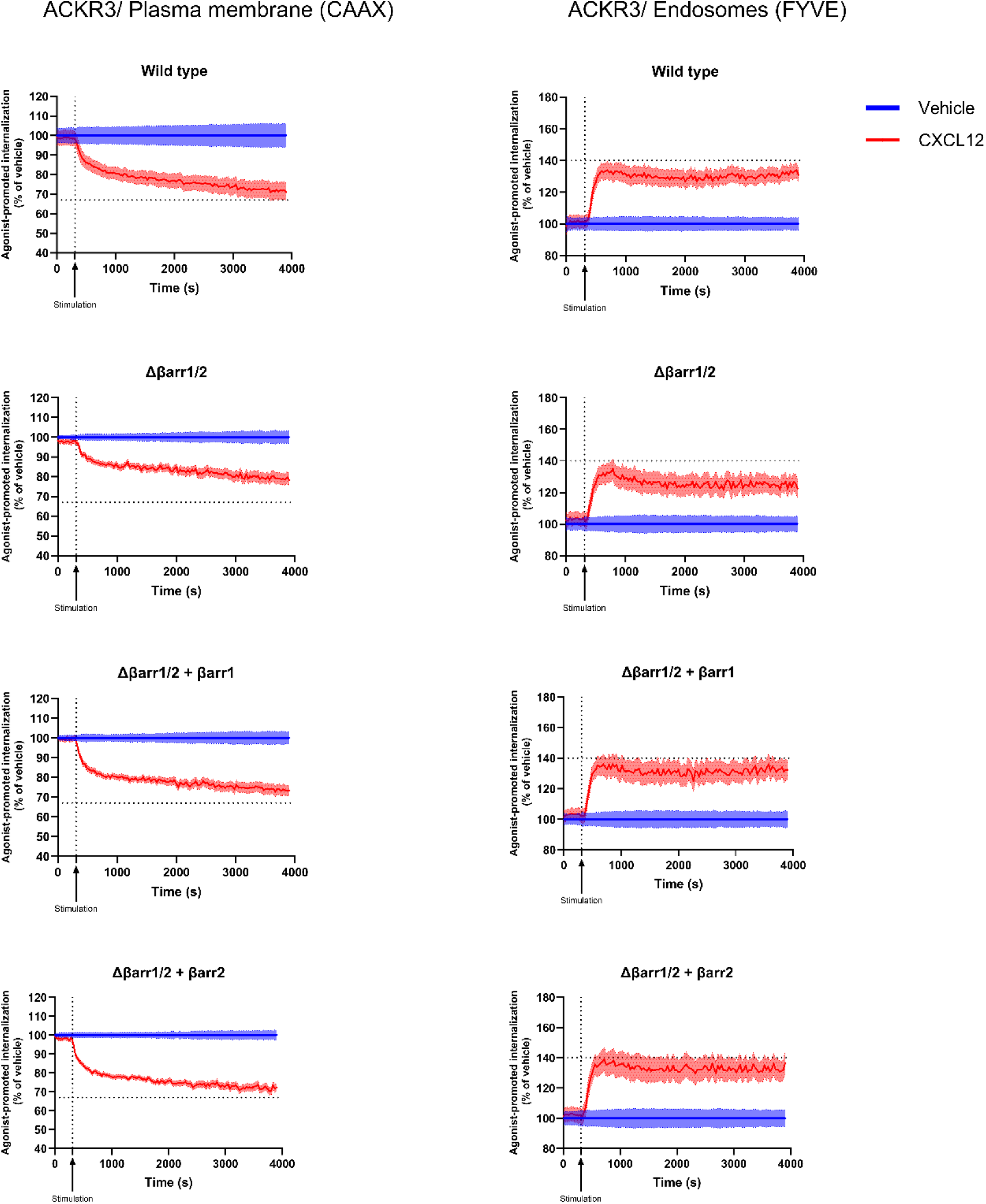
Intracellular trafficking of ACKR3 in HEK293 cells in the presence and absence of β-arrestins. Ligand induced internalization and early endosome association of ACKR3 were assessed in HEK293 cells (wild type) or depleted for β-arrestin isoforms (Δβarr1/2) transfected or not with plasmids encoding the indicated β-arrestins (βarr1 or βarr2). Cells were co-transfected with plasmids encoding ACKR3-RLucII and rGFP-CAAX (left panels, CAAX) or rGFP-FYVE (right panels, FYVE). Cells were incubated in the absence (vehicle) or presence of ligand (CXCL12 100 nM). Means ± SEM of triplicates from three independent experiments. The BRET signal was recorded in kinetic mode for 60 min, with ligand stimulation applied at 5 min, as indicated by the arrow (stimulation). The BRET signal was normalized to the vehicle control, which was set to 100%. The maximal response observed in wild-type cells is represented by a horizontal line and was used as a reference for β-arrestin depleted cells (Δβarr1/2) as well as β-arrestin rescue cells (Δβarr1/2 + βarr), allowing direct comparison across conditions. All data are presented as mean ± SEM of triplicates from three independent experiments

**Table 1.**
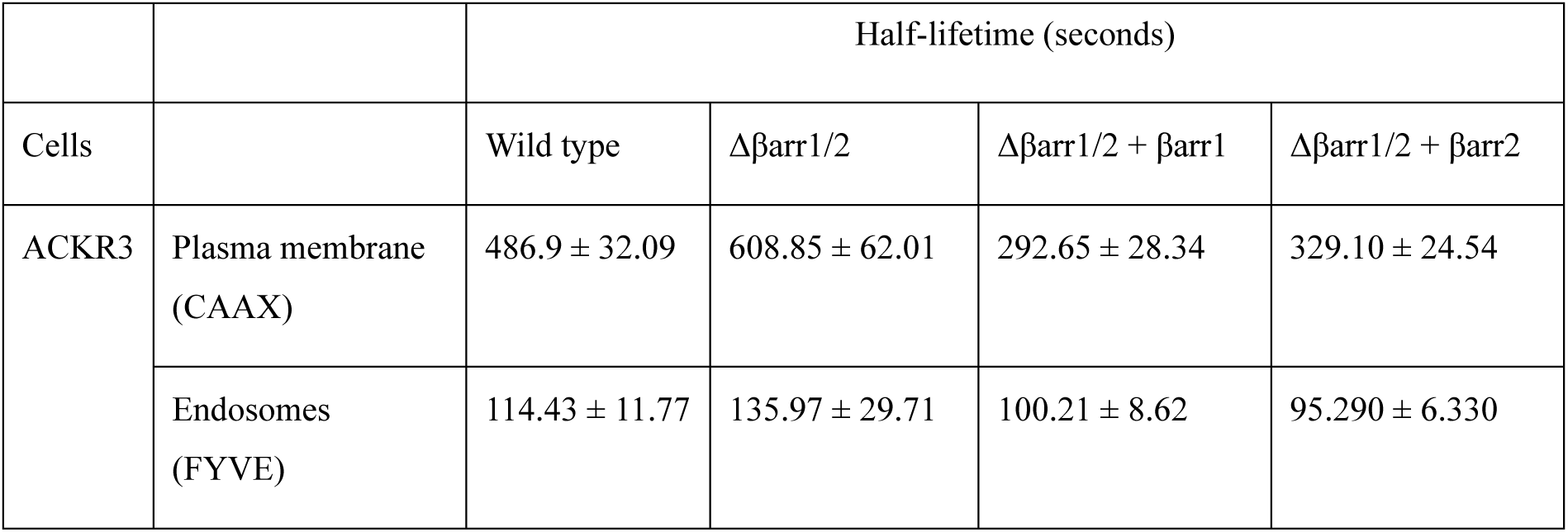
Half-lifetime of ACKR3 internalization kinetics in the presence and absence of β-arrestins. The kinetics of ligand-induced plasma membrane dissociation (CAAX) and early endosome association (FYVE) of ACKR3 were measured in HEK293 wild-type cells or in cells depleted of β-arrestin isoforms (Δβarr1/2) and reconstituted with either β-arrestin1 (βarr1) or β-arrestin2 (βarr2). Kinetic measurements were performed as shown in Figure 3. The half-lifetime values of receptor internalization under the different conditions are reported here. Data represent the mean ± SEM of triplicates from three independent experiments.

As shown in Supplementary Figure 4, The EC_50_ for both CXCL12 and CXCL11-12 to promote the loss of receptor from the plasma membrane and the association with the endosomes (loss of cell surface receptor: CXCL12 = 0.36 ± 0.45nM, CXCL11-12 = 0.31 ± 0.18nM; endosome translocation: CXCL12 = 1.12 ± 0.10nM, CXCL11-12 = 1.48 ± 0.14nM are in good agreement with the reported binding affinities of CXCL12 and CXCL11-12 on ACKR3 (CXCL12: between 0.4 and 1.3nM; CXCL11-12: between 0.1 and 1nM) [13, 14, 48].

### Dynamics of ACKR3-β-arrestin interactions

Given that we found that β-arrestins are dispensable for ACKR3 endocytosis, we wanted to reassess the recruitment of β-arrestins under the same conditions in the same cell backgrounds. Here we quantitatively assessed the ability of the two chemokines CXCL12 and CXCL11-12 to promote β-arrestin1 and β-arrestin2 recruitment to the plasma membrane. HEK293 (Figure 4) and HeLa (Supplementary Figure 5) cells were transfected with rGFP-CAAX, ACKR3 and either β-arrestin1-RlucII or β-arrestin2-RlucII. Stimulation for 10 min with CXCL12 or CXCL11-12 induced a concentration-dependent increase of the BRET for both β-arrestin isoforms in the two cell types. The data indicate that both chemokines promote a robust β-arrestin recruitment in either HEK293 or HeLa cells even though this recruitment is not necessary for receptor endocytosis.

**Figure 4.**
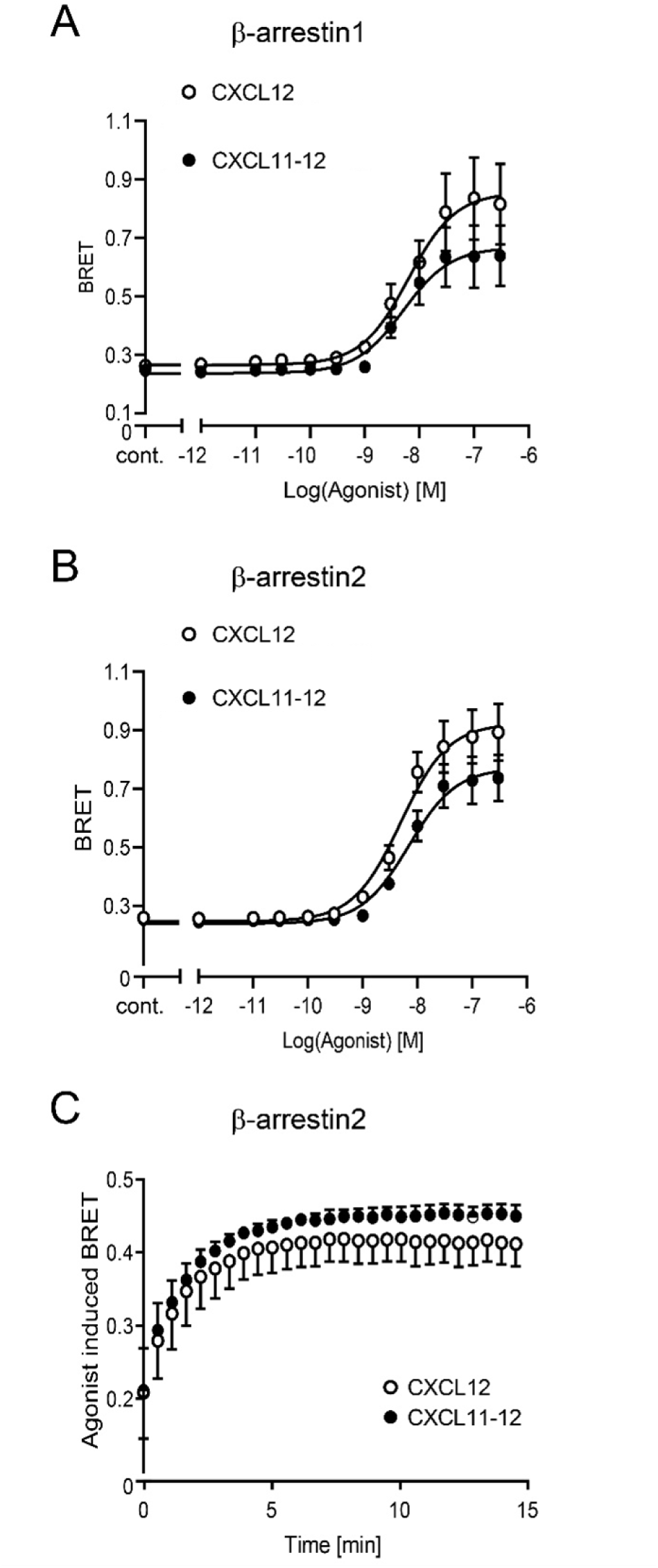
Agonist-dependent recruitment of β-arrestins to the plasma membrane in HEK293 cells. HEK293 cells were transfected with plasmids encoding ACKR3 together with rGFP-CAAX and β-arrestin1-RlucII (**A**) or β-arrestin2-RlucII (**B**). Cells were stimulated for 10 min with the indicated concentrations of CXCL12 or the ACKR3 selective ligand CXCL11-12 while BRET signals were recorded. Ligand induced both β-arrestins recruitment to ACKR3 with EC50 for β-arrestin1 (CXL12 = 6.01 ± 0.19 nM; CXCL11-12 = 5.07 ± 0.20 nM) and for β-arrestin2 (CXL12 = 4.95 ± 0.12 nM; CXCL11-12 = 6.94 ± 0.12 nM). Data are presented as mean ± SEM of duplicates from six independent experiments. (C) Kinetics of ligand-induced β-arrestin2 recruitment to the plasma membrane in HEK293 cells transfected with plasmids encoding ACKR3, rGFP-CAAX and β-arrestin2-RlucII. Cells were stimulated for 15 min with 50 nM of CXCL12 or CXCL11-12. BRET signals were recorded at 30 second intervals. Data are presented as mean ± SEM of triplicates from three independent experiments.

Ligand-induced chemokine receptor internalization is known to depend on the GTPase dynamin which pinches off clathrin-coated nascent endosomes from the plasma membrane [54]. To determine whether ACKR3 endocytosis, which occurs independently of β-arrestins, requires dynamin, we used a dominant-negative dynamin mutant (K44A) lacking GTPase activity [55]. As shown in Supplementary Figure 6, expression of K44A-dynamin completely blocked chemokine mediated ACKR3 endocytosis and scavenging, indicating that ACKR3 internalizes through a dynamin-dependent pathway despite being β-arrestin independent.

### Role of ACKR3 C-terminus phosphorylation in chemokine scavenging and receptor trafficking

Given that we observed concomitant chemokine-promoted β-arrestin recruitment and ACKR3 endocytosis and given that β-arrestins had previously been implicated in ACKR3 endocytosis and chemokine scavenging, we aimed at further assessing the potential role of β-arrestin in this process. For this purpose, we took advantage of previous observations indicating that replacing all nine serine and threonine residues (S335, T338, T341, S347, S350, T352, T355, S360 and T361) to alanine in the C-terminus of ACKR3 (ACKR3-ST/A) abolished both constitutive and ligand induced β-arrestin2 recruitment [21, 45, 56]. First, we confirmed that in contrast to HEK293 cells expressing wild-type (WT) ACKR3-YFP stimulation of cells expressing ACKR3-ST/A-YFP did not promote the recruitment of β-arrestin2 as assessed by BRET between receptor-YFP and β-arrestin2-RlucII (Figure 5A). Consistently, β-arrestin2 recruitment to ACKR3 was completely abolished in HEK293 lacking all GRKs constitutively expressed in the cells (ΔGRK2/3/5/6), whereas re-introduction of either GRK2 or GRK5 restored β-arrestin2 recruitment (Figure 5B).

**Figure 5.**
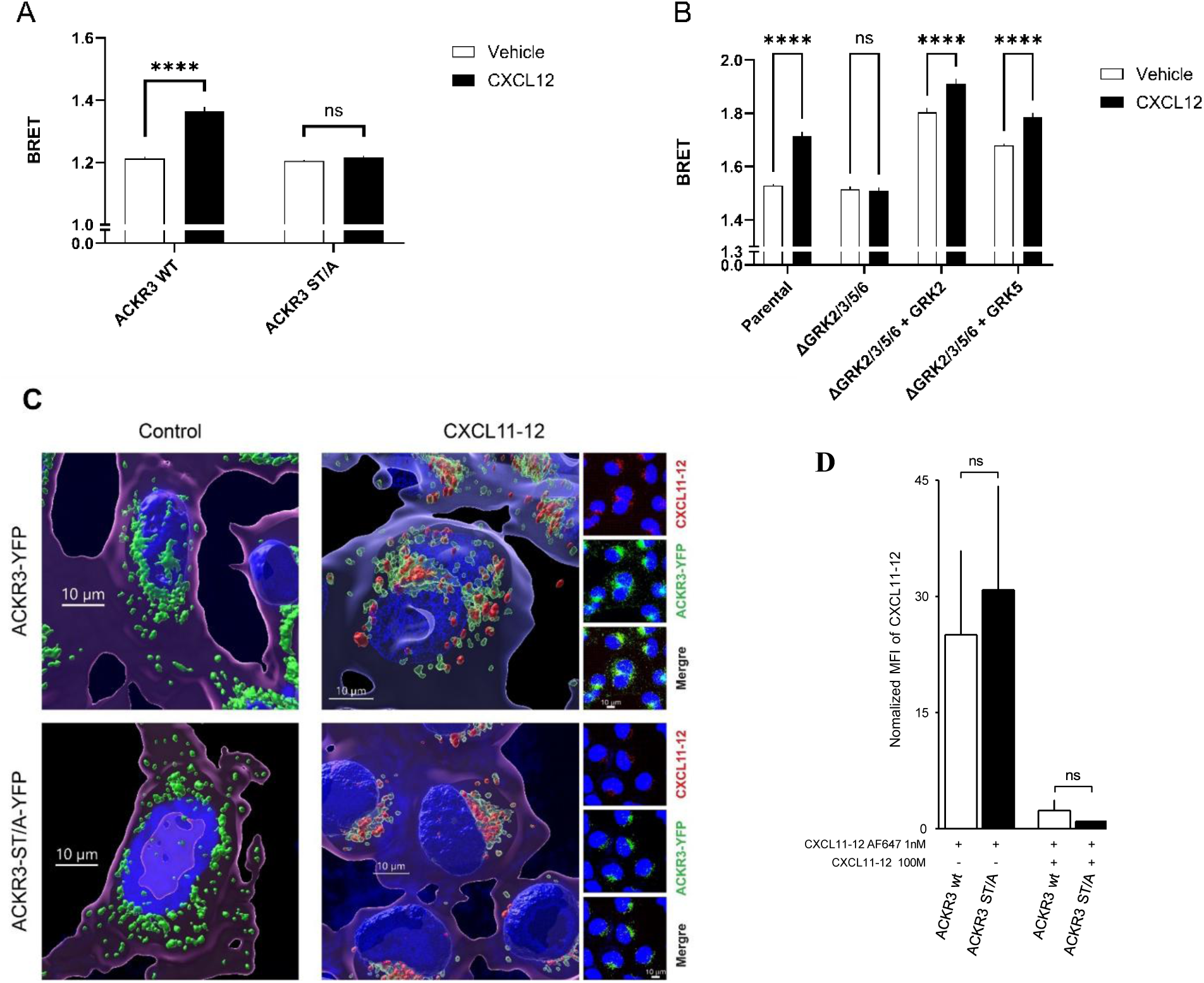
Phosphorylation deficient ACKR3 scavenges chemokines without β-arrestin recruitment. **(A)** Recruitment of β-arrestin2 to the wild-type (WT) or phosphodeficient mutant (ST/A) of ACKR3 was assessed in HEK293 cells transfected with plasmids encoding human ACKR3-wt-YFP or ACKR3-ST/A-YFP and β-arrestin2-RlucII. Black bars: cells were stimulated with 100 nM CXCL12 for 20 min at 37°C while BRET signals were recorded. Data are presented as mean ± SEM of triplicates from four independent experiments. Statistical analysis was performed using two tailed unpaired T-test; **** p < 0.0001, ns: not significant. (B) Recruitment of β-arrestin2 to ACKR3 wild-type was assessed in HEK293 cells (parental) or depleted for GRK2, GRK3, GRK5, GRK6 (ΔGRK2/3/5/6) transfected with plasmids encoding the indicated GRK (GRK2 or GRK5) and ACKR3-wt-YFP with β-arrestin2-RlucII. Cells were stimulated with 100 nM CXCL12 for 20 min at 37°C while BRET signals were recorded. Data are presented as mean ± SEM of triplicates from three independent experiments. Statistical analysis was performed using two-way ANOVA followed by Sidak’s multiple comparison test (ns: not significant, ****p<0.0001). (C) Confocal imaging of chemokine uptake by wild type and phosphorylation deficient ACKR3-ST/A. HEK293 stably transfected with ACKR3-YFP (upper panels) and ACKR3-ST/A-YFP (lower panels) were imaged by confocal microscopy either untreated (left panels, control) or incubated for 30 min at 37°C in the presence of 50 nM CXCL11-12 (right panels, CXCL11-12). Large panels: 3-D representation from Z-stacks (nuclei blue, ACKR3-YFP and ACKR3-(ST/A)-YFP green, CXCL11-12-AF647 red). Blurred cell surfaces were calculated from the dim signal of plasma membrane-associated YFP using low surface area detail level (1-2 µm) in IMARIS. The purple color surrounding the cells reflect the weak receptor expression at the plasma membrane. The small panels on the right show collapsed stacks, which were used for 3D rendering. (D) Flow cytometry of chemokine uptake by wild-type and phosphorylation deficient ACKR3-ST/A. HEK293 stably transfected with ACKR3-YFP (open bars) and ACKR3-ST/A-YFP (black bars) were incubated with 1 nM CXCL11-12AF647 for 15 min in the absence and presence of 100 nM unlabeled CXCL11-12. Mean fluorescence intensities (MFI) of internalized CXCL11-12AF647 were normalized for receptor expression levels deduced from the YFP signals. Data from three independent experiments, means ± SD. Statistical analysis was performed using two tailed unpaired T-test, ns not significant.

Confocal imaging of HEK293 cells showed that under steady state at 37°C both WT ACKR3-YFP and ACKR3-ST/A-YFP displayed weak surface expression and markedly populated intracellular endosome-like structures (reflecting constitutive endocytosis as seen in Figure 1) regardless of whether the receptor harbors potential phosphorylation sites or not (Figure 5C, left panels). The findings are consistent with our observation that β-arrestins are dispensable for receptor internalization and suggest that phosphorylation of the receptor C-terminus is also unnecessary for its constitutive endocytosis. To corroborate the phosphorylation-independent receptor trafficking, cells expressing either WT- or ST/A- ACKR3-YFP were incubated with fluorescently labeled CXCL11-12 (CXCL11-12-AF647) for 30 min at 37°C. Confocal microscopy revealed that the chimeric chemokine was internalized by both receptor constructs, indicating phosphorylation-independent chemokine scavenging activity of ACKR3 (Figure 5C, right panels). CXCL11-12-AF647 uptake was fully prevented in the presence of a saturating concentration of unlabeled CXC11-12, confirming the selectivity of the scavenging (Figure 5D).

Using BRET sensors to monitor ACKR3 endocytosis upon CXCL11-12 and CXCL12 stimulation, we found that ACKR3-ST/A endocytosis was slower and reduced compared to WT-ACKR3, suggesting that the C-terminal phosphorylation plays a role but is not essential for receptor endocytosis (Figure 6A). This is consistent with the findings of Saaber *et al*. [21] reporting that, in primary neuronal cultures from mice expressing ACKR3-ST/A, receptor internalization and chemokine scavenging were markedly reduced. As observed for ACKR3 endocytosis in the absence of β-arrestins (Figure 3), the impact of phosphorylation site mutations on endocytic processes was more pronounced when considering the loss of cell-surface receptors than endosomal translocation (Figure 6A). This finding is consistent with a role for receptor phosphorylation and β-arrestin recruitment in regulating the kinetics of endocytosis rather than being strictly required for the process. This observation is further supported by the scavenging confocal images (Figure 5C), which reflects the combined contribution of constitutive and agonist-induced mechanisms.

**Figure 6.**
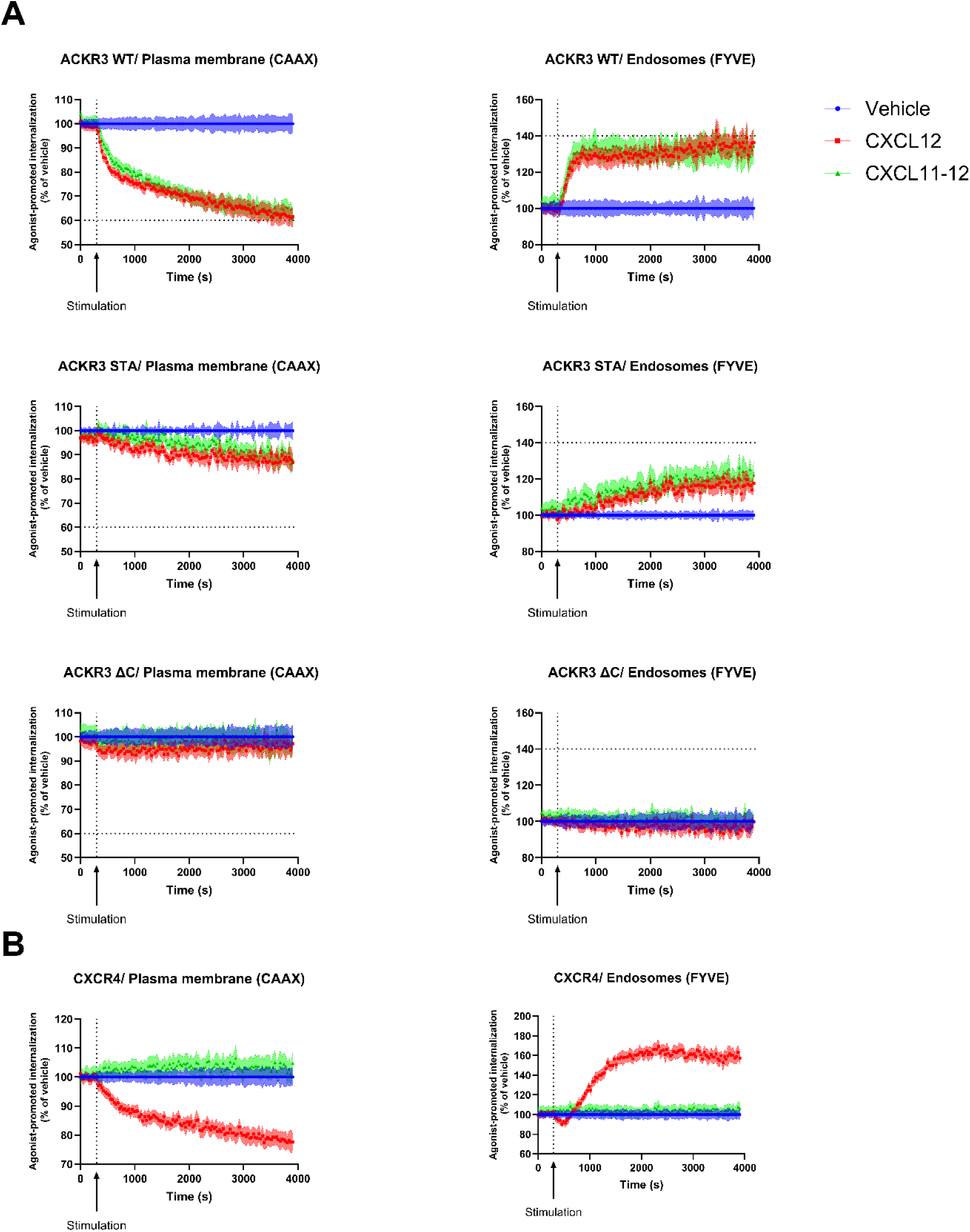
Ligand-induced ACKR3 (WT/STA/ΔC) or CXCR4 endocytosis. **(A)** Ligand induced internalization and early endosome association of ACKR3 were assessed in HEK293 cells transfected with plasmids encoding wild-type (WT) or phosphodeficient mutant (ST/A) or deleted C-terminus (ΔC) ACKR3-RLucII and rGFP-CAAX (left panels, CAAX) or rGFP-FYVE (right panels, FYVE). Cells were incubated in the absence (Vehicle) or presence of 100 nM of ligand (CXCL12 or CXCL11-12). The BRET signal was recorded in kinetic mode for 60 min, with ligand stimulation applied at 5 min, as indicated by the arrow (stimulation). The BRET signal was normalized to the vehicle control, which was set to 100%. The maximal response of wild-type ACKR3 is indicated by a horizontal line, which is also applied to the ACKR3 mutants (ST/A and ΔC) to enable direct comparison. (B) Ligand induced internalization and early endosome association of CXCR4 were assessed in HEK293 cells transfected with plasmids encoding wild-type CXCR4-RLucII and rGFP-CAAX (left panel, CAAX) or rGFP-FYVE (right panel, FYVE). Cells were incubated in the absence (Vehicle) or presence of 100 nM of ligand (CXCL12 or CXCL11-12). The BRET signal was recorded in kinetic mode for 60 min, with ligand stimulation applied at 5 min, as indicated by the arrow (stimulation). The BRET signal was normalized to the vehicle control, which was set to 100%. All data are presented as mean ± SEM of triplicates from three independent experiments.

Contrasting with the remaining endocytosis observed for ACKR3-ST/A, a deletion of the entire receptor C-terminus (ACKR3-ΔC) completely abrogated the endocytosis (Figure 6A). This is consistent with a previous reports showing that the C-terminus of ACKR3 is required for spontaneous receptor internalization and chemokine scavenging [15, 50, 57] and suggest that it may play a role that is independent of its phosphorylation. It is noteworthy that the endocytosis promoted by CXCL11-12 cannot result from CXCR4 activation since as shown in Figure 6B, CXCl1-12 in contrast to CXCL12 cannot promote the endocytosis of CXCR4.

### Role of GRK2 and GRK5 in ligand-dependent ACKR3 trafficking

Given the fact that ACKR3 trafficking may occur independently of its C-terminus phosphorylation, we asked whether GRKs are necessary. For this purpose, we measured ACKR3 trafficking in HEK293 ΔGRK2/3/5/6 cells complemented or not with plasmids expressing GRK2, GRK5 or both. In contrast to what is observed in WT cells, no ACKR3 internalization was observed in the absence of GRK2/3/5/6 upon stimulation with CXCL12 or CXCL11-12. Reintroduction of GRK2 and GRK5 individually or together rescued the agonist-promoted ACKR3 endocytosis (Figure 7A). Although Schafer, *et al*. suggested that the role of GRK2 in ACKR3 endocytosis resulted from the activation of endogenous CXCR4 by CXCL12 [40], this cannot explain the robust effect of GRK2 observed in our study since CXCL11-12 does not activate CXCR4 and does not promote its endocytosis (Figure 7B). The role of GRK2 did not simply result from its over-expression since significant CXCL12- and CXCL11-12-promoted ACKR3 endocytosis could also be observed in HEK293 cells lacking either GRK5/6 (ΔGRK5/6) or GRK2/3 (ΔGRK2/3) (Supplementary Figure 7A) confirming that endogenously expressed GRK2/3 or GRK5/6 can promote endocytosis. As expected, only CXCL12 and not CXCL11-12 promoted the endocytosis of CXCR4 in these cells (Supplementary Figure 7B).

**Figure 7.**
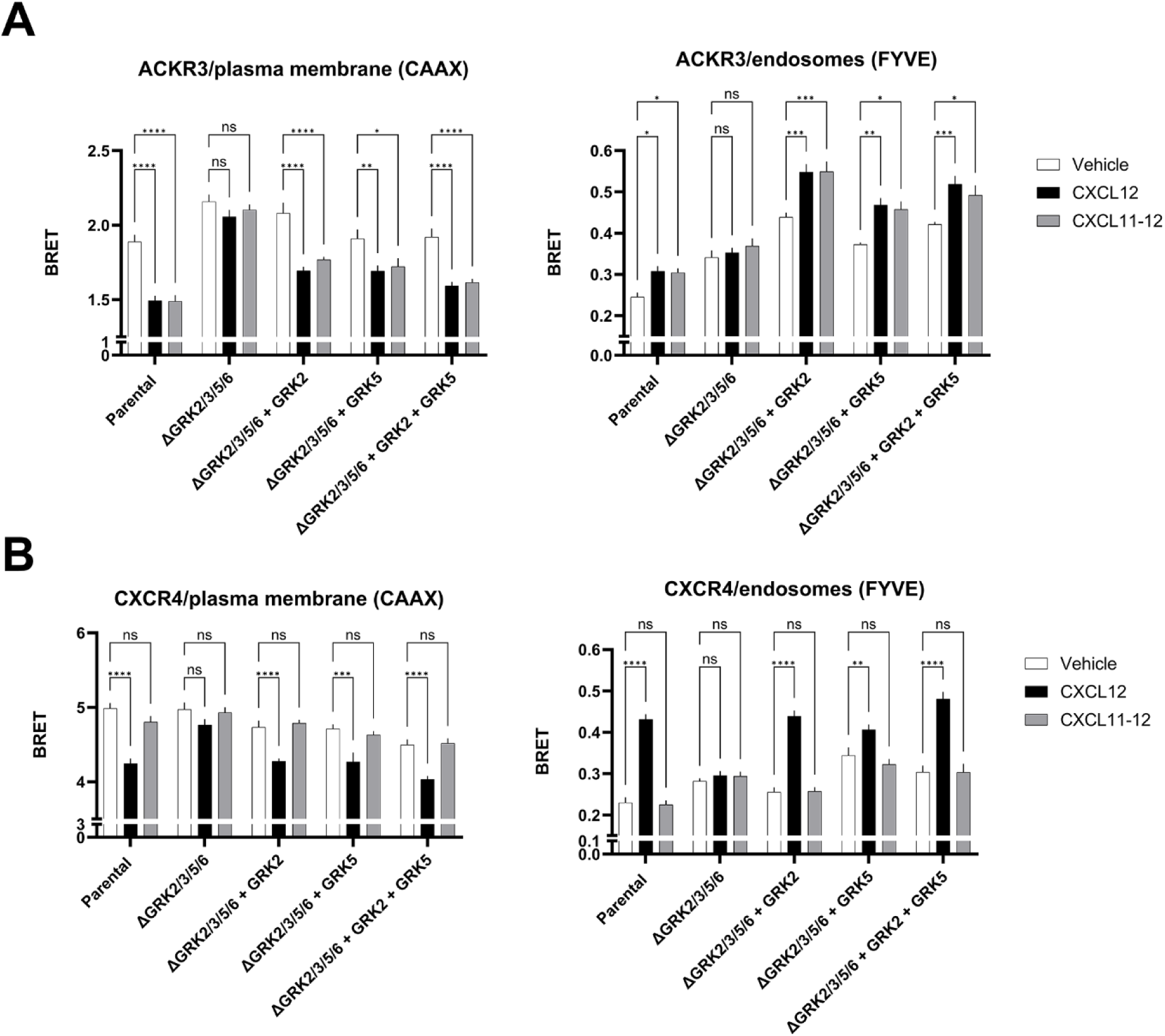
Ligand-dependent ACKR3 and CXCR4 trafficking in the presence or absence of GRK2 and GRK5. **A.** Ligand-induced internalization and early endosome association of ACKR3 and CXCR4 were assessed in parental HEK293 cells or in cells depleted of GRK2, GRK3, GRK5, and GRK6 (ΔGRK2/3/5/6), transfected with plasmids encoding the indicated GRKs (GRK2, GRK5, or both). Cells were co-transfected with plasmids encoding ACKR3-RLucII (**A**) or CXCR4-RLuc (**B**) and either rGFP-CAAX (left panel, CAAX) or rGFP-FYVE (right panel, FYVE). Cells were incubated for 25 min in the absence of ligand (white bars) or in the presence of 100 nM CXCL12 (black bars) or 100 nM CXCL11-12 (grey bars). Data are presented as mean ± SEM of triplicates from three independent experiments. Statistical analysis was performed using two-way ANOVA followed by Tukey’s multiple comparison test (ns, not significant; *p < 0.05; **p < 0.01; ***p < 0.001; ****p < 0.0001).

### GRK2 recruitment to ACKR3 upon chemokine activation

To ascertain that GRK2 directly promotes ACKR3 endocytosis, we monitored its recruitment to ACKR3 by BRET measuring the interaction between GRK2–GFP10 and ACKR3-RlucII. Stimulation with either CXCL12 or CXCL11-12 induced a recruitment of GRK2 to ACKR3, similar to that observed for CXCR4 upon activation by CXCL12 (Figure 8). As expected, stimulation with CXCL11-12 did not promote GRK2 recruitment to CXCR4, confirming the selectivity of CXCL11-12 promoted GRK2 recruitment to ACKR3. Consistent with the observation that chemokines-promoted endocytosis of the phosphorylation-deficient ACKR3 mutant form (ACKR3-ST/A) (Figure 6A) and the proposed role of GRK2 in the endocytic process (Figures 7A and Supplementary Figure 7A, GRK2 can be directly recruited to ACKR3-ST/A as detected by BRET (Figure 8). The rationale that led investigators to propose that GRK5/6 and not GRK2/3 should be recruited to ACKR3 is based on the fact that free Gβγ released from activated G protein promotes the recruitment of GRK2 but is not required for GRK5/6 [58] and that ACKR3 does not activate any G proteins [27]. To assess whether GRK2 promoted endocytosis required free Gβγ, we used a GRK2 fragment tethered to the plasma membrane via a CAAX motif (known as βARKct-CAAX) that sequesters free Gβγ subunits [59]. As expected, βARKct-CAAX completely blocked the GRK2- mediated CXCR4 endocytosis in ΔGRK2/3/5/6 cells (Figure 9B). βARKct-CAAX was also found to block GRK2-promoted MOR endocytosis upon met-enkephalin stimulation (Figure 9C), consistent with the reported role of Gβγ for GPCRs that can activate G proteins. In contrast, the Gβγ scavenger had no impact on the ACKR3 endocytosis promoted by GRK2 upon stimulation with either CXCL12 or CXCL11-12 (Figure 9A) indicating that free Gβγ is not essential for the action of GRK2 on ACKR3. Similarly, βARKct-CAAX had no effect on ACKR3 endocytosis observed in ΔGRK5/6 (Supplementary Figure 8) confirming that GRK2/3 promoted ACKR3 endocytosis does not require G protein activation and free Gβγ release. Consistent with the lack of free Gβγ requirement for ACKR3 endocytosis, a GRK2 construct lacking the PH domain (GRK2 ΔPH) required for GRK2 recruitment by Gβγ [58] was able to restore ACKR3 endocytosis in ΔGRK2/3/5/6 cells but not the endocytosis of CXCR4 or MOR that require Gβγ for GRK2 recruitment (Figure 10).

**Figure 8.**
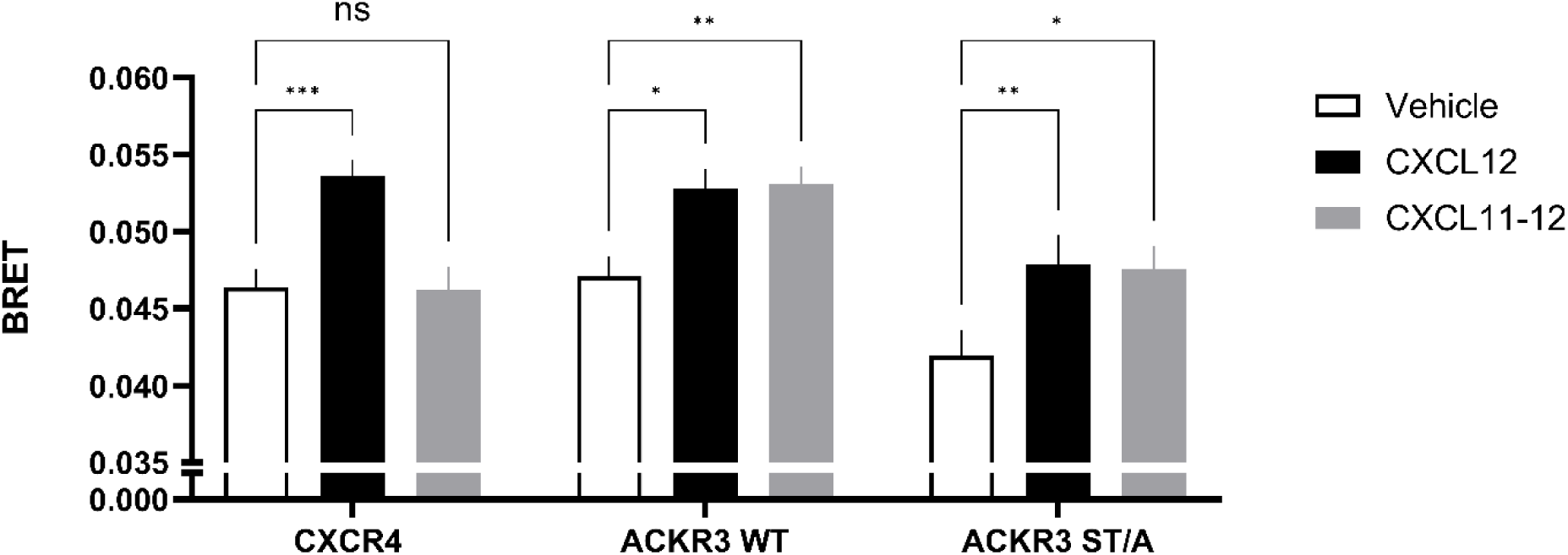
Ligand-induced GRK2 recruitment following ACKR3 or CXCR4 stimulation. Ligand-induced GRK2 recruitment was measured in HEK293 cells co-transfected with plasmids encoding CXCR4-RLucII or ACKR3-RLucII, either wild-type (WT) or the phosphodeficient mutant (ST/A), together with GRK2-GFP10. Cells were incubated for 2 min in the absence of ligand (white bars) or in the presence of 100 nM CXCL12 (black bars) or 100 nM CXCL11-12 (grey bars). Data are presented as the mean ± SEM of triplicates from three independent experiments. Statistical analysis was performed using two-way ANOVA followed by Dunnett’s multiple comparison test (ns, not significant; *p < 0.05; **p < 0.01; ***p < 0.001).

**Figure 9.**
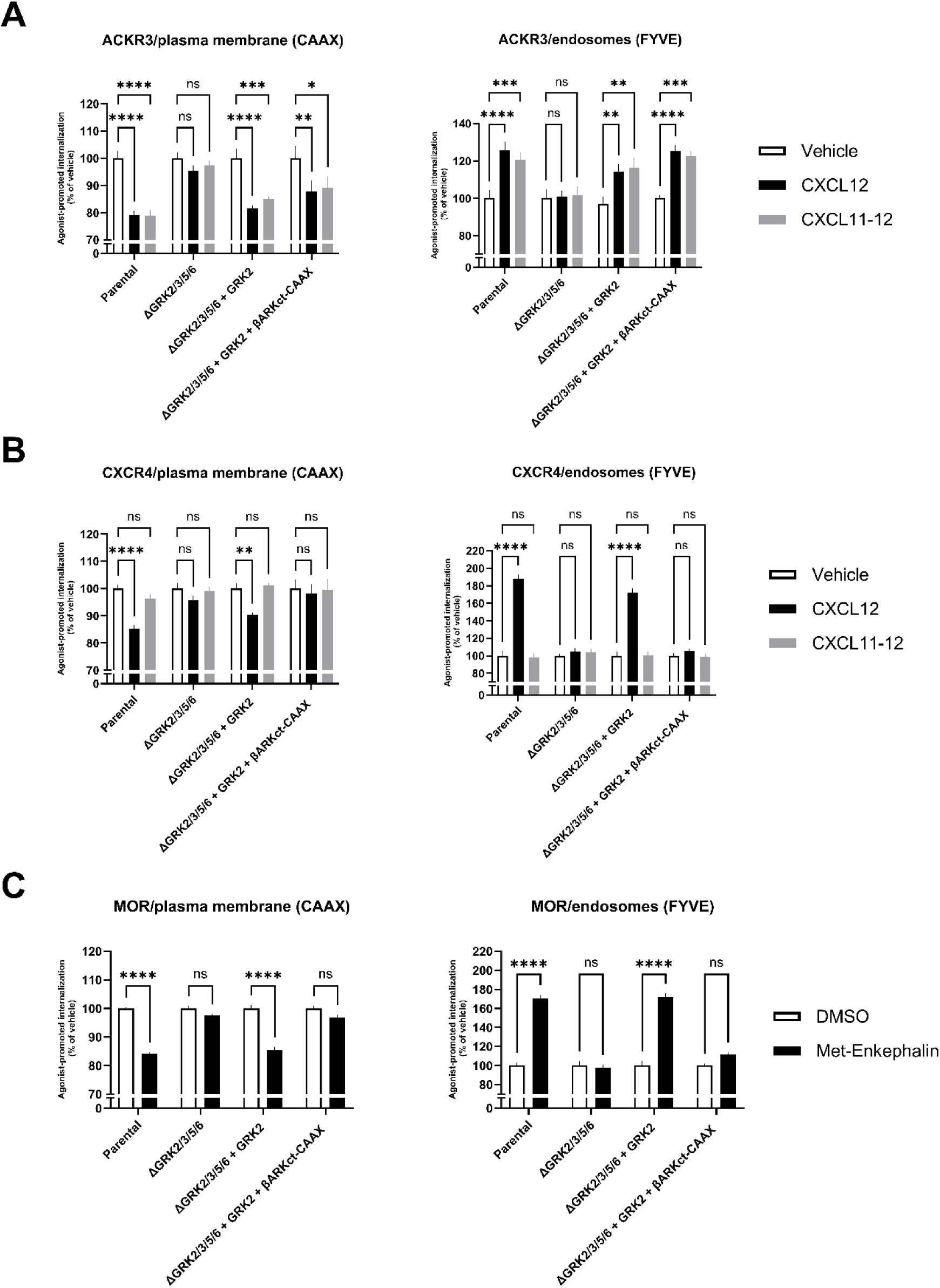
GRK2-dependent ligand-induced trafficking of ACKR3, CXCR4, and MOR in the presence or absence of Gβγ or Gβγ blocker (βARKct-CAAX). Ligand-induced internalization and early endosome association of ACKR3, CXCR4 and MOR were assessed in parental HEK293 cells or in cells depleted of GRK2, GRK3, GRK5, and GRK6 (ΔGRK2/3/5/6), transfected or not with a plasmid encoding GRK2 in the presence or absence of a plasmid encoding βARKct-CAAX. Cells were co-transfected with plasmids encoding ACKR3-RLucII (**A**), CXCR4-RLuc (**B**) and MOR-RLuc (**C**) and either rGFP-CAAX (left panel, CAAX) or rGFP-FYVE (right panel, FYVE). Cells were incubated for 25 min in the absence of ligand (white bars, **A**, **B** and **C**), or in the presence of 100 nM CXCL12 (black bars) or 100 nM CXCL11-12 (grey bars) (**A** and **B**) or in the presence of 5 µM Met-enkephalin (black bars, **C**). Statistical analysis was performed using two-way ANOVA followed by Dunnett’s (**A**) or Tukey’s (**B** and **C**) multiple comparison test (ns, not significant; *p < 0.05; **p < 0.01; ***p < 0.001; ****p < 0.0001). All data are presented as mean ± SEM of triplicates from three independent experiments.

**Figure 10.**
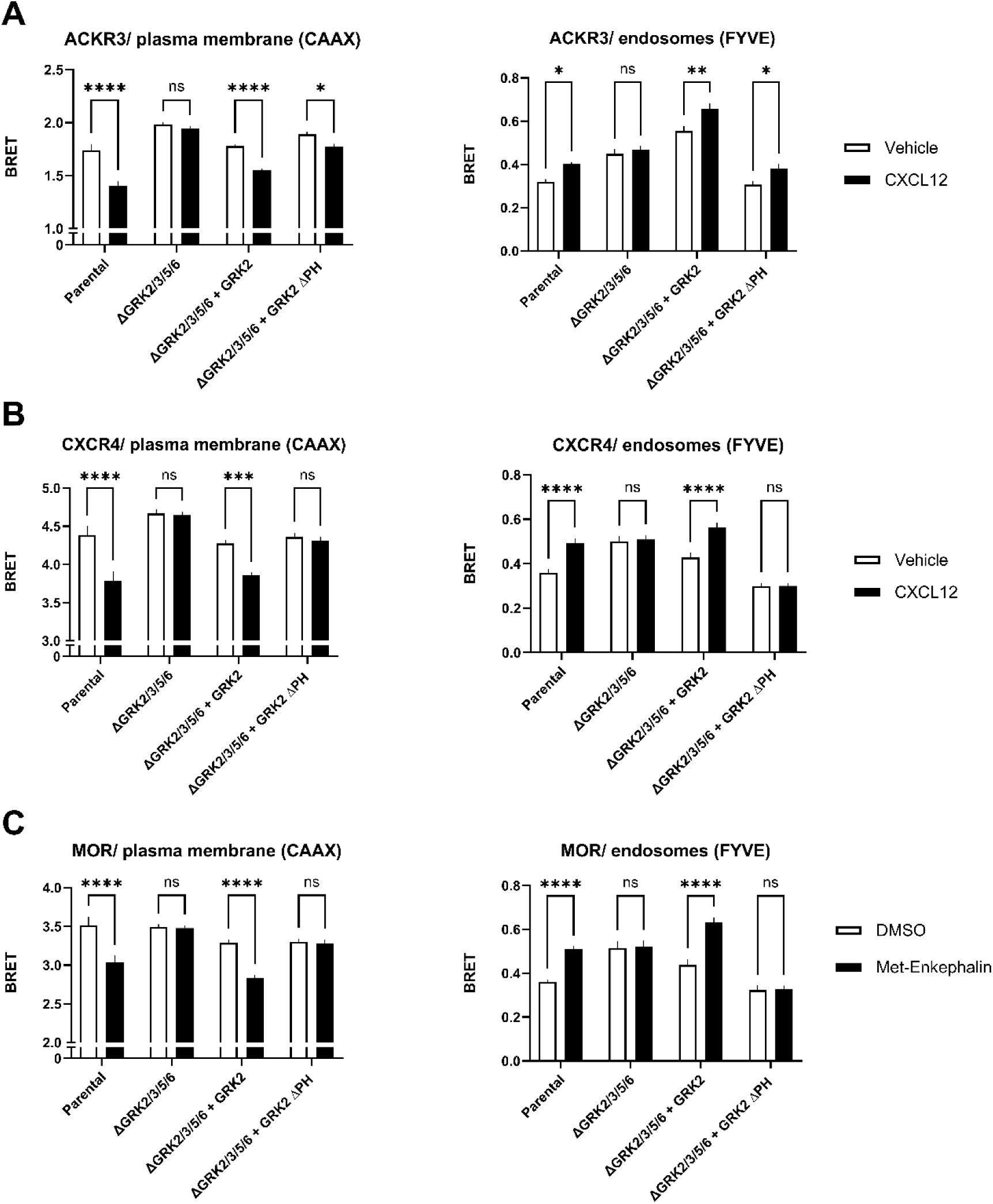
Ligand-induced trafficking of ACKR3, CXCR4, and MOR in the presence or absence of GRK2 or GRK2 lacking PH domain (GRK2 ΔPH). Ligand-induced internalization and early endosome association of ACKR3 (**A**), CXCR4 (**B**) or MOR (**C**) were assessed in parental HEK293 cells or in cells depleted of GRK2, GRK3, GRK5 and GRK6 (ΔGRK2/3/5/6), transfected or not with plasmids encoding GRK2 or GRK2 lacking PH domain (GRK2 ΔPH). Cells were co-transfected with plasmids encoding ACKR3-RLucII (**A),** CXCR4 (**B**) or MOR (**C**) and either rGFP-CAAX (left panel, CAAX) or rGFP-FYVE (right panel, FYVE), and were incubated for 25 min in the absence of ligand (white bars), or in the presence of 100 nM CXCL12 (black bars, **A** and **B**) or of 5 µM Met-enkephalin (black bar, **C**). Statistical analysis was performed using two-way ANOVA followed by Sidak’s multiple comparison test (ns, not significant; *p < 0.05; **p < 0.01; ***p < 0.001; ****p < 0.0001). All data are presented as mean ± SEM of triplicates from three independent experiments.

### Role of GRK2 kinase activity in chemokine promoted ACKR3 endocytosis

As shown previously, ACKR3 phosphorylation is dispensable for receptor internalization, whereas GRK expression is required. We therefore investigated whether the kinase activity of GRK2 contributes to this process. To address this question, we assessed ACKR3 trafficking from the plasma membrane to endosomes in HEK293 ΔGRK2/3/5/6 cells complemented with plasmids encoding kinase-dead mutants of GRK2 (GRK2 K220R) [60]. As shown in Figure 11A, GRK2 K220R restored ACKR3 internalization in ΔGRK2/3/5/6 cells lacking endogenous GRKs. To verify that these GRK mutants were indeed catalytically inactive, CXCR4 and MOR whose internalization strictly depend on GRK kinase activity were used as a control. As expected, GRK2 K220R failed to rescue the endocytosis of CXCR4 (Figure 11B) and MOR (Supplementary Figure 9) in ΔGRK2/3/5/6 cells, confirming the loss of kinase activity. Further supporting the observation that phosphorylation of ACKR3 is not required for its GRK2-dependent endocytosis, chemokine-promoted endocytosis of ACKR3 ST/A in ΔGRK2/3/5/6 cells could be restored by re-expression of GRK2 (Supplementary Figure 10).

**Figure 11.**
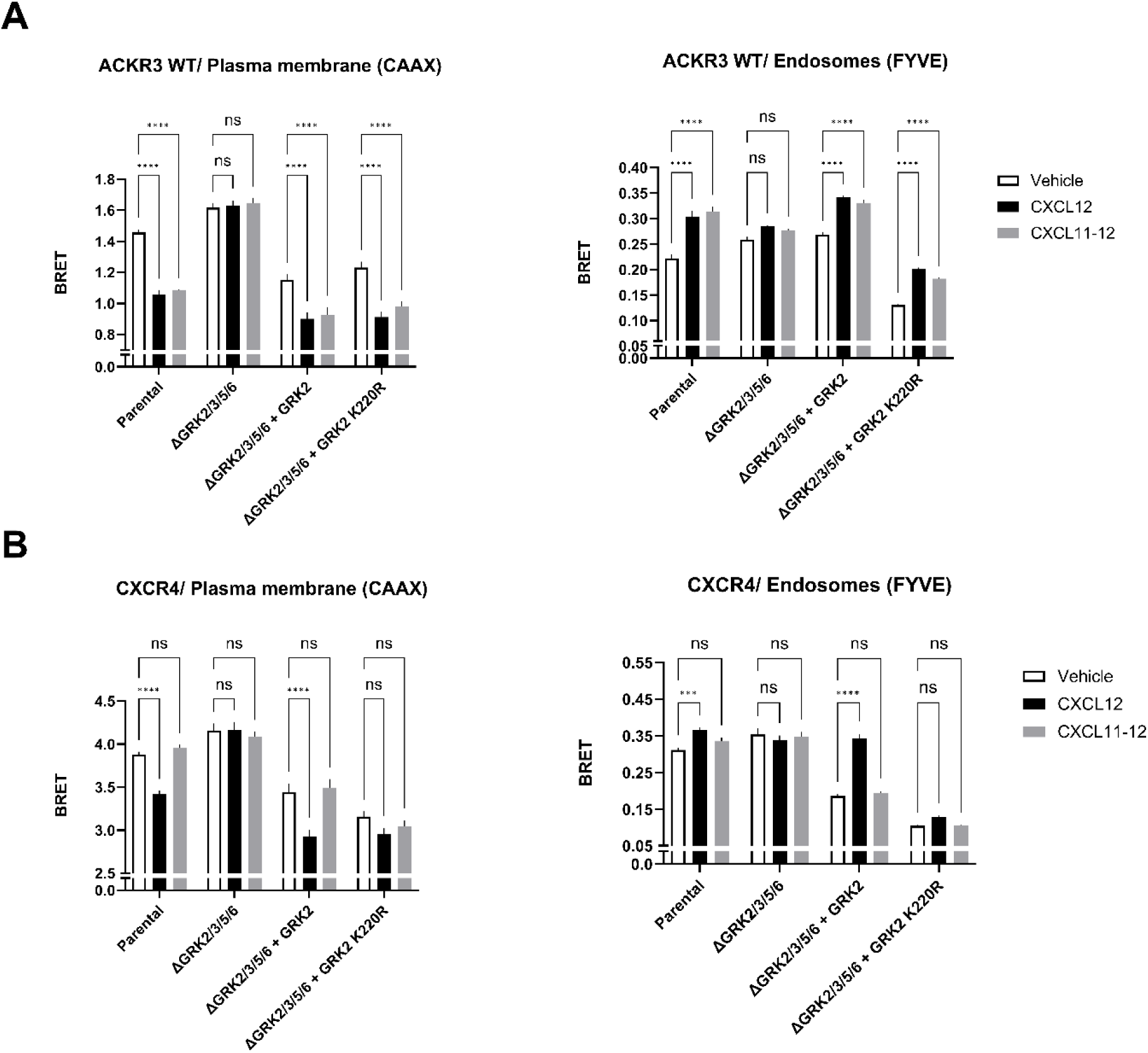
Ligand-dependent ACKR3 and CXCR4 trafficking in the presence or absence of kinase-dead GRK2 (K220R). Ligand-induced internalization and early endosome association of ACKR3 or CXCR4 were assessed in parental HEK293 cells or in cells depleted of GRK2, GRK3, GRK5, and GRK6 (ΔGRK2/3/5/6), transfected or not with plasmids encoding GRK2 or GRK2 K220R (kinase-dead mutant). Cells were co-transfected with plasmids encoding ACKR3-RLucII (**A**) or CXCR4-RLuc (**B**) and either rGFP-CAAX (left panel, CAAX) or rGFP-FYVE (right panel, FYVE). Cells were incubated for 25 min in the absence of ligand (white bars) or in the presence of 100 nM CXCL12 (black bars) or 100 nM CXCL11-12 (grey bars). Data are presented as mean ± SEM of triplicates from three independent experiments. Statistical analysis was performed using two-way ANOVA followed by Dunnett’s multiple comparison test (ns, not significant; ***p < 0.001; ****p < 0.0001).

## Discussion

ACKR3 is frequently upregulated in pathological circumstances. In cancer, the over-expression of ACKR3 has been associated with poor outcome [61–65]. Targeting ACKR3 pharmacologically is therefore of potential interest for the treatment of debilitating diseases. However, a better understanding of the receptor mode of action is necessary to fully understand its role in oncogenesis. Pharmacological inhibition of ACKR3 leads to 2-3-fold increased serum levels of CXCL12 in human and mouse [66–68], demonstrating the importance of its scavenging activity in chemokine homeostasis.

The notion that ACKR3 does not couple to G proteins and the observation of spontaneous as well as ligand-promoted ACKR3 endocytosis [15] suggested that ACKR3 is naturally biased towards β-arrestins [27, 29], a protein classically know to be involved in GPCR endocytosis, and that such coupling must be responsible for the receptor’s scavenging activity. In fact, β-arrestin2 recruitment assays showed that the adaptor is indeed an interactor of ACKR3 [37, 44, 45]. Yet, seemingly conflicting reports have raised questions about the role of β-arrestins and receptor phosphorylation by specific GRKs in ACKR3 endocytosis and chemokine scavenging.

Consistent with a role of β-arrestin in the scavenging activity of the receptor, Luker *et al.* showed that ACKR3-mediated CXCL12 uptake and receptor endocytosis was disrupted in mouse embryonal fibroblasts derived from β-arrestin2 KO but not β-arrestin1 KO mice [36]. Using a β-arrestin dominant-negative mutant and siRNA targeting both β-arrestin1 and β-arrestin2, Canals *et al*. concluded that β-arrestins are required for chemokine-promoted ACKR3 cell surface loss assessed by ELISA and also reported that a mutant form of ACKR3 lacking all phosphorylation sites at the C-terminus was unable to interact with β-arrestins (ACKR3-ST/A) and did not undergo agonist-promoted endocytosis [45]. In sharp contrast, Saaber e*t al.* found that in neuronal cell cultures both β-arrestins were dispensable for receptor chemokine uptake suggesting that they were dispensable for receptor endocytosis. They found, using the ACKR3-ST/A construct, that phosphorylation at the C-terminus of ACKR3 is not essential but contributes to high efficiency chemokine scavenging [21]. Montpas *et al*. for their part reported that siRNA-mediated knock-down of β-arrestin1, β-arrestin2 or both did not affect chemokine degradation and that such degradation was similar in mouse embryonic fibroblasts-derived from wild-type or β-arrestin1/2 KO mice concluding that β-arrestins are dispensable for chemokine scavenging [30]. The lack of β-arrestin involvement in ACKR3 endocytosis and chemokine scavenging was also suggested in HEK293 cells lacking β-arrestins [40]. Regarding possible distinct roles of the two β-arrestins, Saaber *et al.* suggested that although ACKR3 could recruit β-arrestin2 it did not recruit β-arrestin1 in neuronal cells [21]. Conflicting results on the relative roles of different GRKs in ACKR3 trafficking and chemokines were also reported. Using compound 101 as a selective inhibitor of GRK2/3, Saaber *et al.* reported that GRK2/3 mediated phosphorylation plays an important role for the efficient scavenging of chemokines [21]. More recently, using GRK2/3 vs GRK5/6 KO HEK293 cells, Parishmita *et al.* reported that ACKR3 phosphorylation by GRK5/6 but not GRK2/3 was essential for chemokine-promoted β-arrestin recruitment [39]. Similarly, Schafer *et al.* found that deleting GRK5/6 has a major impact on the chemokine-promoted recruitment of β-arrestin2 whereas the deletion GRK2/3 had no effect but that deletion of both GRK5/6 and GRK2/3 had a greater effect than the deletion of GRK5/6 alone [40]. A similar preponderant role of GRK5/6 was found for the chemokine-induced ACKR3 endocytosis and chemokine scavenging [40].

To provide a mechanistic explanation for these seemingly conflicting results on the roles of β-arrestins, GRKs and receptor phosphorylation in both ACKR3 endocytosis and chemokines scavenging we systematically assess the engagement of both β-arrestins by ACKR3, their roles as well as the roles of the GRKs and receptor phosphorylation in the endocytosis of the receptor and chemokine scavenging using orthogonal assays in well controlled systems.

We show here, using confocal microscopy and real-time BRET-based biosensors that both β-arrestins are recruited to ACKR3 upon stimulation with either CXCL12 or the ACKR3 specific chimeric ligand CXCL11-12, and that both β-arrestins are dispensable for ACKR3 endocytosis using two different cell types. Confocal microscopy clearly showed that ACKR3 co-localized with CXCL11-12 in endosomal structures independently of the presence of β-arrestins. Although β-arrestins are fully dispensable for both, ligand-dependent and -independent internalization, BRET sensors revealed that β-arrestins are recruited to the receptor and that the presence of β-arrestins may accelerate the chemokine-promoted internalization consistent with an accessory role. This may partly explain the apparent contradictions about the role of β-arrestins in previous studies.

β-arrestin-independent endocytosis has previously been reported for several GPCRs [60, 69–74]. For the protease-activated receptor-1 (PAR1), β-arrestins are required for receptor phosphorylation-dependent desensitizing uncoupling from G proteins, but not for receptor internalization [70]. For CXCR4, previous studies also reported β-arrestin-independent endocytosis [52, 75], a finding corroborated in our study. As for ACKR3, although the agonist-promoted endocytosis was equivalent in WT and Δβarr1/2, the re-expression of β-arrestins modestly favored endocytosis pointing again to an accessory but not essential role. This is also consistent with the apparent modest contribution of β-arrestins to the constitutive endocytosis observed for ACKR3 in our microscopy studies. The observation that, in contrast to most GPCRs, β-arrestins are recruited to ACKR3 but do not play an essential role in its endocytosis raises questions about the possible distinct conformation that β-arrestin adopts upon binding to receptors. For instance, it has been proposed that recruitment of β-arrestin as a result of distinct receptor phosphorylation patterns may have different outcomes on desensitization, endocytosis and downstream signaling [24, 26, 76, 77]. The independence from β-arrestin for ACKR3 endocytosis and chemokine scavenging role also raise questions about an alternative role for the scaffolding protein. Given that β-arrestins have been suggested as a scaffolding protein involved in the activation of ERK, it is noteworthy that ACKR3 has been proposed to promote the activation of ERK1/2 and K-Ras in a β-arrestin-dependent manner [78].

Scavenging of CXCL12 and ACKR3 endocytosis has previously been shown to strongly depend on an intact C-terminus of the receptor [15, 45, 50, 56]. The requirement of the C-terminus for ACKR3 endocytosis was confirmed in the present study. Taken with the efficient spontaneous and ligand-promoted internalization of ACKR3 in the absence of β-arrestins these data points towards an alternative mechanism for ACKR3 internalization that involves the C-terminus of the receptor. The most prominent post-translational modification of GPCRs C-termini is phosphorylation [24, 26, 45, 79]. Here using a mutant form of ACKR3 lacking all the C-terminus phosphorylation sites, ACKR3-ST/A, we found that, although phosphorylation clearly accelerates and increase the extent of endocytosis, the phosphorylation sites were found to be dispensable as ACKR3-ST/A undergoes robust endocytosis. When considering the scavenging activity, we found that the ACKR3-ST/A could still efficiently scavenge CXCL11-12 suggesting that the reduced level of endocytosis observed for this phosphorylation-deficient mutant is sufficient to support scavenging. This is consistent with the observation of Saaber *et al*. who found that the ST/A mutation attenuated but did not abolish scavenging. It is noteworthy that expression of ACKR3-ST/A prevented the perinatal death observed in in the ACKR3 KO mice [21] indicating that phosphorylation is not essential for the critical physiological role of ACKR3 for organismal survival.

Intriguingly, although phosphorylation is dispensable, we found that the presence of GRK is essential for the receptor endocytosis as no agonist-promoted endocytosis was observed in GRK2/3/5/6 KO cells. This is consistent with a similar observation made by Schafer *et al.* [40]. In their studies, comparing ΔGRK2/3 with ΔGRK5/6 the authors suggested that GRK5/6 has a predominant role with GRK2/3 having only a marginal role. In contrast, we found that either GRK2 or GRK5 re-expression restored similar endocytosis levels. A confounding factor when determining the roles of specific GRKs using KO cells is that the endogenous levels of the different kinases vary across cell types. For instance, GRK2/3 is notoriously expressed at very low levels in HEK293 cells compared to GRK5/6 that could explain the relatively little effect of deleting or knocking-down GRK2/3 in these cells. In contrast, the higher level of GRK2/3 in neurons [80] could explain why, in neuronal cells, Saaber *et al.* found a preponderant role for GRK2/3 [21].

Yet, the role of GRK2/3 is surprising given the lack of coupling of ACKR3 to G proteins [27] and the role of Gβγ in promoting the translocation of GRK2/3 to the plasma membrane [58]. In their study, Schafer suggests that the Gβγ could be provided through the activation of endogenously expressed CXCR4 by CXCL12. This cannot be argued to explain our results since CXCL11-12 is selective for ACKR3 and does not activate CXCR4. Consistent with a direct role of GRK2 on ACKR3 endocytosis independently of G protein activation, its effect was found to be independent of Gβγ, as it could not be blocked with the Gβγ scavenger βARKct, and that a GRK2 mutant form lacking the Gβγ binding PH domain could also restore ACKR3 endocytosis. Notably, GRK2 recruitment to GPCRs in the absence of G protein activation has also been described for other receptors, including the dopamine D2 receptor [81, 82] and the β₂-adrenergic receptor [83], further supporting our conclusions. These data suggest that, at least for some receptors, the affinity of the agonist-bound receptor for GRK2 may be sufficient to promote its recruitment without the additional contribution of Gβγ.

The fact that phosphorylation of the receptor’s C-terminus is not essential for endocytosis but that GRKs are, is counter-intuitive and suggests that the kinase activity is not essential for its role in endocytosis. In agreement with this notion, we found that a kinase-dead form of GRK2 could restore chemokine-promoted endocytosis of ACKR3 in ΔGRK2/3/5/6 cells. Together, these results support a model in which GRK2 can promote ACKR3 endocytosis in a phosphorylation-independent manner, although phosphorylation may accelerate or enhance the process, consistent with a scaffolding role for GRKs. To explore how GRK2 could interact with ACKR3, an AlphaFold3 prediction of an ACKR3-GRK2 complex was generated using the AlphaFold server [84]. As shown in Figure 12, the N-terminus of GRK2 can directly interact in the pocket created by the opening of the receptor’sTM6 which normally forms the interaction site for the C-terminal α5 helix of G proteins for receptors that can couple to G proteins [85]. Similar structural Cryo-EM GRK-GPCR complexes showing this N-terminal engagement of GRKs in the receptor were reported for the rhodopsin-GRK1 [86] and the NTSR1-GRK2 [87] complexes suggesting that the mechanism described here for ACKR3 could be more general.

**Figure 12.**
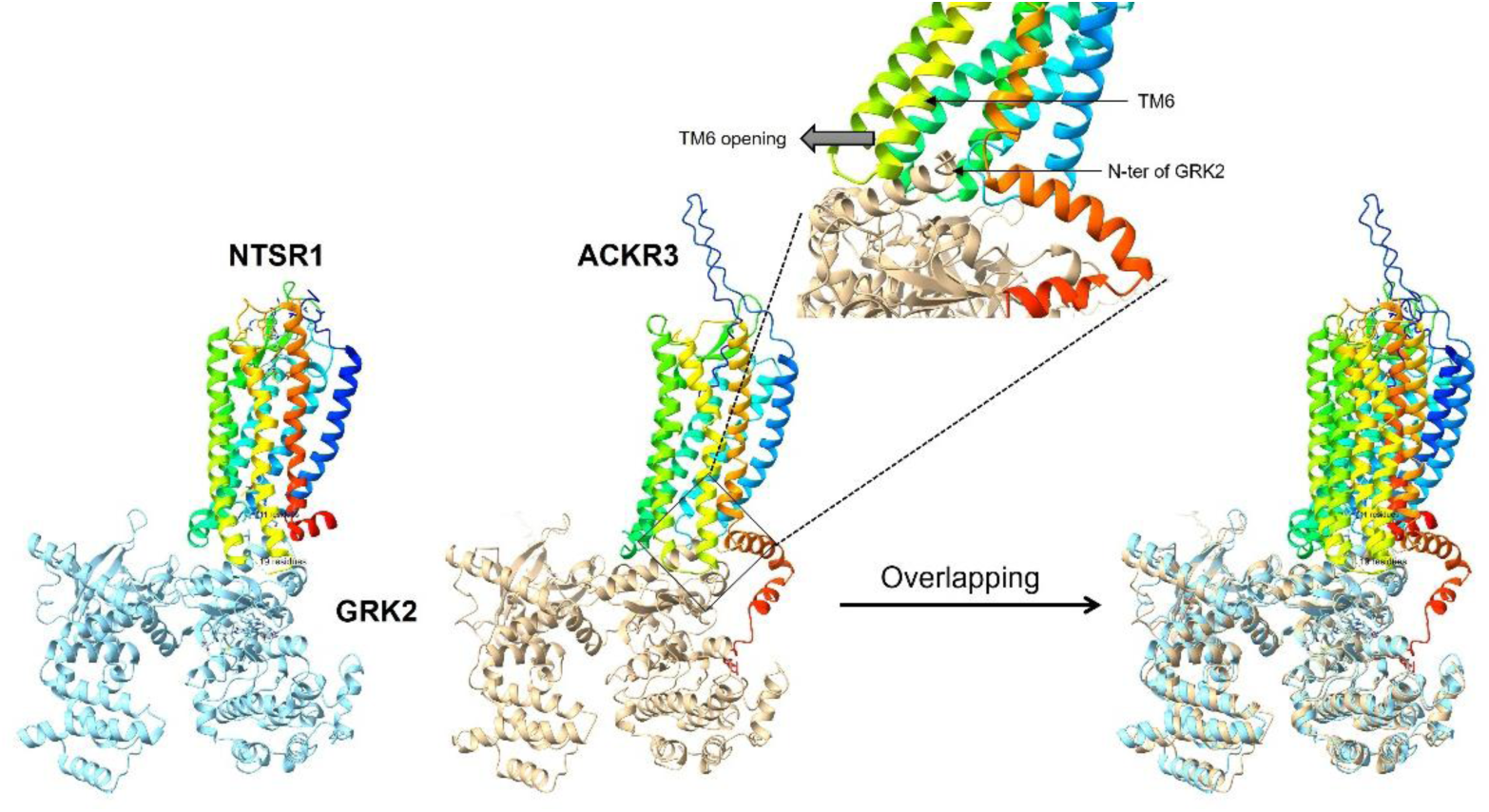
AlphaFold3 prediction of the ACKR3–GRK2 complex. The amino acid sequences of human ACKR3 and GRK2 were retrieved from the UniProt database (accession numbers P25106 and P25098, respectively). For the AlphaFold3 prediction of the ACKR3–GRK2 complex, the sequences were submitted to the AlphaFold3 server for structure prediction. The server generated a compressed output file containing predicted structural models ranked by confidence and energy score. The model with the lowest predicted energy was selected, and the corresponding CIF file was opened and visualized using ChimeraX. For comparison, the cryo-EM structure of the NTSR1–GRK2–Gαq complex (PDB ID: 8JPB) was downloaded from the Protein Data Bank and opened in ChimeraX. The Gαq subunit was removed from the structure. The NTSR1–GRK2 and ACKR3–GRK2 models were then rotated and aligned to obtain a similar orientation and subsequently superimposed. Images show NTSR1–GRK2 (left), ACKR3–GRK2 (middle), and the superimposed models (right). NTSR1 and ACKR3 are displayed in rainbow colors, while GRK2 is shown in blue (NTSR1 complex) and brown (ACKR3 complex). The zoomed and rotated view (top) highlights the outward movement of ACKR3 transmembrane helix 6 (TM6) and the insertion of the GRK2 N-terminus into the intracellular cavity.

Taken together our results reconcile some of the apparently contradictory data of the literature and show that although β-arrestin and receptor phosphorylation may facilitate ACKR3 endocytosis and chemokine scavenging, they are clearly dispensable. Yet, GRKs are essential for this process most-likely as a scaffolding protein that does not rely only on its kinase activity. The mechanism by which GRKs can directly supports endocytosis remains to be investigated.

## Material and Methods

### Plasmids

The plasmids encoding pcDNA3.1-PCP-ACKR3 [50], ACKR3-RlucII [88], ACKR3ΔC-RlucII [89], CXCR4-RlucII [52], MOR-RlucII [90], V2R-RlucII [52], rGFP-CAAX and rGFP-FYVE [51], β-arrestin 1 or 2 [52], β-arrestin 1- or 2-RlucII [52], dynamin K44A [52, 91], GRK2, GRK5 and GRK2 K220R [92], βARKct-CAAX [52, 59], and GRK2-GFP10 [90] have been previously described. GRK2 lacking the PH domain (GRK2ΔPH) was generated using GRK2-GFP10 as a template. The GRK2 sequence lacking the PH domain (amino acids 558 to 652) was amplified by PCR and subcloned into pCDNA3.1 digested with NheI and BamHI, followed by ligation using T4 DNA ligase.

Mouse ACKR3-ST/A was PCR amplified from spleenocytes isolated from an ACKR3-ST/A mouse [21], cloned into pCDH-EF1-MCS-COP-GFP (Adgene) and the sequence verified. Human ACKR3-ST/A was generated by standard PCR using proof reading TAQ-polymerase and the mouse sequence as template. The reverse primer included the sequence of the human ACKR3 extreme C-terminus (the only difference between the human mouse sequence is at amino acid position 360 (hu: Ser; mouse: Asn). To generate ACKR3-YFP fusion contracts the receptor genes were fused with YFP inserting a linker (LESGGGGSGGGG SGGGGS) upstream of the YFP gene and cloned into the pALPS lentivector as transfer plasmid [93]. pcDNA3.1 constructs encoding wild-type and ST/A ACKR3-YFP were also generated for BRET assays. These inserts were isolated by digesting pALPS-ACKR3-YFP with XbaI and BamHI, then subcloned into pCDNA3.1 digested with NheI and BamHI, followed by ligation using T4 DNA ligase.

### Reagents

Recombinant CXCL12 and CXCL11-12 were expressed, purified and, when indicates, labeled as previously described [48, 94] or purchased from R&D Systems. Met-enkephalin was from Cayman Chemical Company (Item No. 23284). Lipofectamine™ LTX Reagent with PLUS™ Reagent, Vybrant™ DiOCell-Labeling solutions as well as all culture media and supplements were from Thermo Fisher. Polyethylenimine (PEI) was from Polysciences. SFP-Synthase (phosphopantetheinyl transferase) was prepared following the manufacturer’s instruction (Covalys Biosience). CoA-Atto dyes (Atto565 and Atto647N, Atto-Tec) were prepared as described [94]. Poly-D-lysine coated glass bottom dishes were from MatTek Corporation. Tyrode’s buffer (140 mM NaCl, 2.7 mM KCl, 1 mM CaCl_2_, 12 mM NaHCO_3_, 5.6 mM D-glucose, 0.5 mM MgCl_2_, 0.37 mM NaH_2_PO_4_, 25 mM Na-HEPES pH 7.4) was prepared in house. Prolume Purple and Coelentrazine H were purchased from Nanolight Technology.

### Cell lines

HEK293SL (referred throughout as HEK293 cells) were selected in Laporte’s laboratory (McGill University) [51, 95]. Deletion of β-arrestin1 and 2 in HEK293 and HeLa cells were previously described [23, 46]. In HeLa and HEK293 cells, β-arrestin1 and 2 were deleted by sequential CRISPR/Cas9 methodology using the following commercial plasmids as described [46]. Cells were transfected using Lipofectamine LTX or polyethylenimine (PEI), according to the manufacturers’ instructions, for microscopy and BRET assays, respectively. All cell lines were grown in Dulbecco’s Modified Eagle’s medium (DMEM) supplemented with 10% fetal bovine serum and 1% w/v penicillin–streptomycin at 37°C in a humidified atmosphere at 95% air and 5% CO_2_. All cell culture reagents were from Thermo Fisher. HEK293 cells with quadruple GRK knockout (ΔGRK2/3/5/6) as well as double knockouts (ΔGRK2/3 and ΔGRK5/6) were provided by Asuka Inoue (Tohoku University) and have been previously described [96].

### Transfection

48 hours prior to BRET measurements, HEK293 cells were transfected using PEI with the indicated biosensors (1 µg total DNA per 3.5 × 10⁵ cells). Cells were seeded in poly-D-lysine–coated 96-well plates at a density of 35,000 cells per well. For reconstitution of β-arrestin expression in Δβarr1/2 cells, 200 ng of plasmid encoding either β-arrestin1 or β-arrestin2 was transfected per 3.5 × 10⁵ cells. For GRK expression in ΔGRK2/3/5/6 cells, cells were transfected with 50 ng of GRK2 (wild type or mutants) or 5 ng of GRK5 per 3.5 × 10⁵ cells. For Gβγ scavenging, ΔGRK2/3/5/6 and ΔGRK5/6 cells were transfected with 500ng of plasmid encoding βARKct-CAAX per 3.5 × 10⁵ cells.

For confocal imaging, HeLa cells were seeded on 35mm glass bottom dishes. After 24 hours, cells were transfected with the indicated plasmids using Lipofectamine LTX (for each plate: 0.4 µg of DNA, 0.4 µl of Plus reagent, 1.5 µl of Lipofectamine LTX).

### Confocal microscopy and PPTase labeling

HeLa cells were plated as above at a density of 5 x 10^4^ cells per dish. After 24 hours, cells were transfected with 400 ng of PCP-ACKR3 alone or 100 ng of PCP-ACKR3 + 300 ng of rGFP-CAAX or rGFP-FYVE. HEK293 cells (10^6^) were nucleofected with 2.5 µg of PCP-ACKR3 + 2.5 µg of rGFP-CAAX or rGFP-FYVE. 24 hours post plasmid application, cells were washed with Dulbecco’s modified phosphate-buffered saline (DPBS) supplemented with 10 mM MgCl_2_ (DPBSM) and incubated for 30 min at 17°C with 2 μM PPTase and 5 μM CoA-Atto 647N in DPBSM [50]. Cells were rinsed with cold DPBSM and then either immediately fixed with formaldehyde using 4% paraformaldehyde (PFA) or incubated at 37°C for the indicated times in the presence or absence of 50 nM of CXCL11-12 Atto565 [48]. When indicated 5 μM Vybrant™ DiO Cell-Labeling Solution was added. Cells were imaged with a laser scanning confocal microscope (Leica SP5) at high magnification (63x or 100x). Cell center Z-stack planes (3-6) were recorded at 0.2-0.4 µm distance and collapsed for presentation.

### BRET measurements

Cells were co-transfected with a BRET donor and a BRET acceptor per 3.5 × 10⁵ cells. BRET donors included wild-type or ST/A mutant ACKR3-RlucII and CXCR4-RlucII (16 ng for endocytosis assays and 50 ng for GRK2 recruitment assays), ACKR3ΔC-RlucII (32 ng), MOR-RlucII (8 ng), V2R-RlucII (5 ng), and β-arrestin 1- or β-arrestin 2-RlucII (5 ng). BRET acceptors included rGFP-FYVE or rGFP-CAAX (300 ng), ACKR3-YFP (600 ng), and GRK2-GFP10 (400 ng). After 48 hours, cells were washed with PBS, then incubated 1h in Tyrode’s buffer. Cells were then treated with the indicated concentrations of CXCL12, CXCL11-12, Met-enkephalin and AVP in Tyrode’s buffer for different times at 37 °C. Prior to signal acquisition, the substrate Prolume Purple (methoxy e-CTZ; NanoLight Technology, cat. no. 369, final concentration 1 µM) was added and incubated for 15 min for BRET2 measurements (donor: RlucII; acceptor: rGFP or GFP10). For BRET1 measurements (donor: RlucII; acceptor: YFP), coelenterazine h (CTZh; NanoLight Technology, cat. no. 301, final concentration 2.5 µM) was added, followed by incubation for 15 min at 37°C. BRET signals were recorded using either a Mithras™ LB940 multimode microplate reader (Berthold Technologies, Bad Wildbad, Germany) or a Spark multimode reader (Tecan), equipped with appropriate donor and acceptor emission filters. For BRET2, emissions were collected using a donor filter centered at 400 nm (± 70 nm) and an acceptor filter centered at 515 nm (± 20 nm). BRET2 values were calculated as the ratio of GFP emission (515 nm) to RlucII emission (400 nm). Data acquisition was performed using MicroWin 2000 software (Berthold Technologies). For BRET1 measurements, luminescence was detected using emission filters of 430–485 nm for the donor (RlucII) and 505–590 nm for the acceptor (YFP). Data acquisition was performed using Spark Control software. The BRET signal was calculated as the ratio of acceptor emission to donor emission, as previously described [51].

### Data analysis and statistical tests

All quantitative data from BRET and confocal microscopy assays were analyzed using GraphPad Software (GraphPad Prism Version 10.6.1, San Diego, California, USA: GraphPad Software; 2024). The software was used to generate all figures (histograms, kinetic curves and dose–response curves). The half-life (t₁/₂) values of ACKR3 kinetic curves were calculated by nonlinear regression. Plasma membrane dissociation (CAAX) data were fitted using a *plateau followed by one-phase decay* model, whereas endosomal translocation (FYVE) data were fitted using a *plateau followed by one-phase association* model. Sigmoidal dose–response curves were fitted using the *[Agonist] vs. response — Variable slope (three parameters)* nonlinear regression model, and EC₅₀ values were derived from these fits.

Statistical analyses were performed using GraphPad Prism. Comparisons between two groups were conducted using an unpaired Student’s T-test. For experiments involving three or more groups with one independent variable, one-way ANOVA was used. For experiments involving three or more groups with two independent variables, two-way ANOVA was performed. Multiple comparisons were conducted using the default post hoc tests implemented in GraphPad Prism.

### AlphaFold3 prediction and structural comparison of the ACKR3–GRK2 complex

The amino acid sequences of human ACKR3 (UniProt accession number P25106) and GRK2 (UniProt accession number P25098) were retrieved from the UniProt database. The sequences were submitted to the AlphaFold3 server for prediction of the ACKR3–GRK2 complex structure using default parameters [84]. The server generated a compressed output file containing ranked structural models with associated confidence and predicted energy scores. The model with the lowest predicted energy was selected for subsequent analyses. The corresponding CIF file was visualized using ChimeraX (UCSF) [97].

For structural comparison, the cryo-electron microscopy structure of the NTSR1–GRK2–Gαq complex [87] was downloaded from the Protein Data Bank (PDB ID: 8JPB) and opened in ChimeraX. The Gαq subunit was removed prior to analysis. The NTSR1–GRK2 structure and the predicted ACKR3–GRK2 model were manually rotated to obtain comparable orientations and subsequently aligned and superimposed using the matchmaker function in ChimeraX [97]. Structural representations were generated with receptors displayed in rainbow color schemes and GRK2 shown in distinct colors for each complex.

## Supplementary materials

Fig. S1: Plasma membrane staining for quantification.

Fig. S2: Intracellular trafficking of CXCR4 and MOR in HEK293 cells in the presence or absence of β-arrestins.

Fig. S3: Intracellular trafficking of ACKR3 and V2R in HeLa cells in the presence and absence of β-arrestins.

Fig. S4: Agonists dependent internalization of ACKR3 in HEK293 cells.

Fig. S5: Agonist-dependent recruitment of β-arrestins to the plasma membrane in HeLa cells.

Fig. S6: ACKR3 internalization and chemokine scavenging are dynamin dependent.

Fig. S7: Ligand-dependent ACKR3 and CXCR4 trafficking in the presence or absence of endogenous GRK2/3 and GRK5/6.

Fig. S8: Effect of Gβγ blocker (βARKct-CAAX) in GRK2 promoted endocytosis of ACKR3, CXCR4 and MOR in ΔGRK5/6 cells.

Fig. S9: Ligand-dependent MOR trafficking in the presence or absence of kinase-dead GRK2 (K220R).

Fig. S10: Ligand-induced ACKR3 STA endocytosis in the presence or absence of GRK2.

## Acknowledgements

We thank Monique Lagacé for helpful discussion and assistance in manuscript preparation. The authors are thankful to Prs. Stephane Laporte and Asuka Inoue for their generous gift of HEK293SL cells and ΔGRK2/3/5/6, ΔGRK2/3, and ΔGRK5/6 HEK293T cells, respectively. We also thank Prs. Andy Chevigne and Martyna Szpakowska for helpful scientific discussions and valuable suggestions during their visit to M. Bouvier’s laboratory.

## Author contributions

BST, SC, PC MS, BB, ST and PHSP performed experiments, analyzed/interpreted the data; DM performed image analysis; GD and MU provided essential reagents; DL contributed to the experimental strategy; MT and MB conceived the study, supervised the work, interpreted the data, and provided funding. MT, MB, BST, SC and PC wrote the manuscript. All co-authors revised the manuscript.

## Data and materials availability

The Biosensors can be obtained freely for academic research through a regular MTA upon request to MB. All other data needed to evaluate the conclusions in the paper are present in the paper or the Supplementary Materials.

---

**Supplementary Figure 1.**
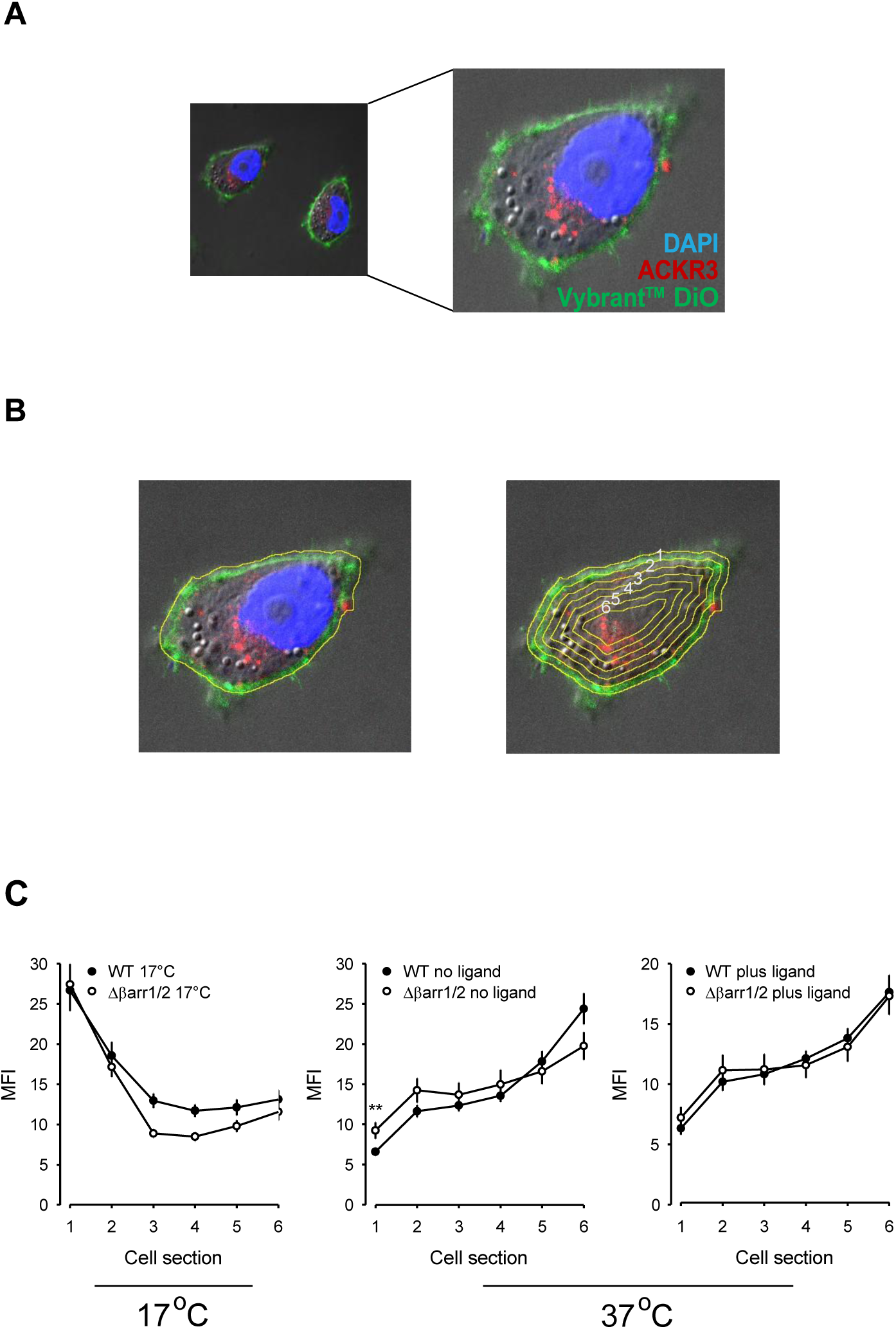
Plasma membrane staining for quantification. **(A)** Plasma membranes of HeLa cells transfected with a plasmid encoding PCP-ACKR3 (red) were stained with Vybrant™ DiO (green) and nuclei with DAPI. (B) The outer region of interest (ROI) was manually drawn corresponding to the plasma membrane (left) and six inner ROI’s were automatically generated at a distance of 1 µm from each other (right). (C) The mean fluorescence intensity (MFI) deriving from labeled ACKR3 was calculated for each ROI with Image J. Means ± SEM from 25-36 cells. Statistical analysis was performed using ordinary one-way ANOVA corrected with Tukey’s multiple comparisons test (**p<0.012).

**Supplementary Figure 2.**
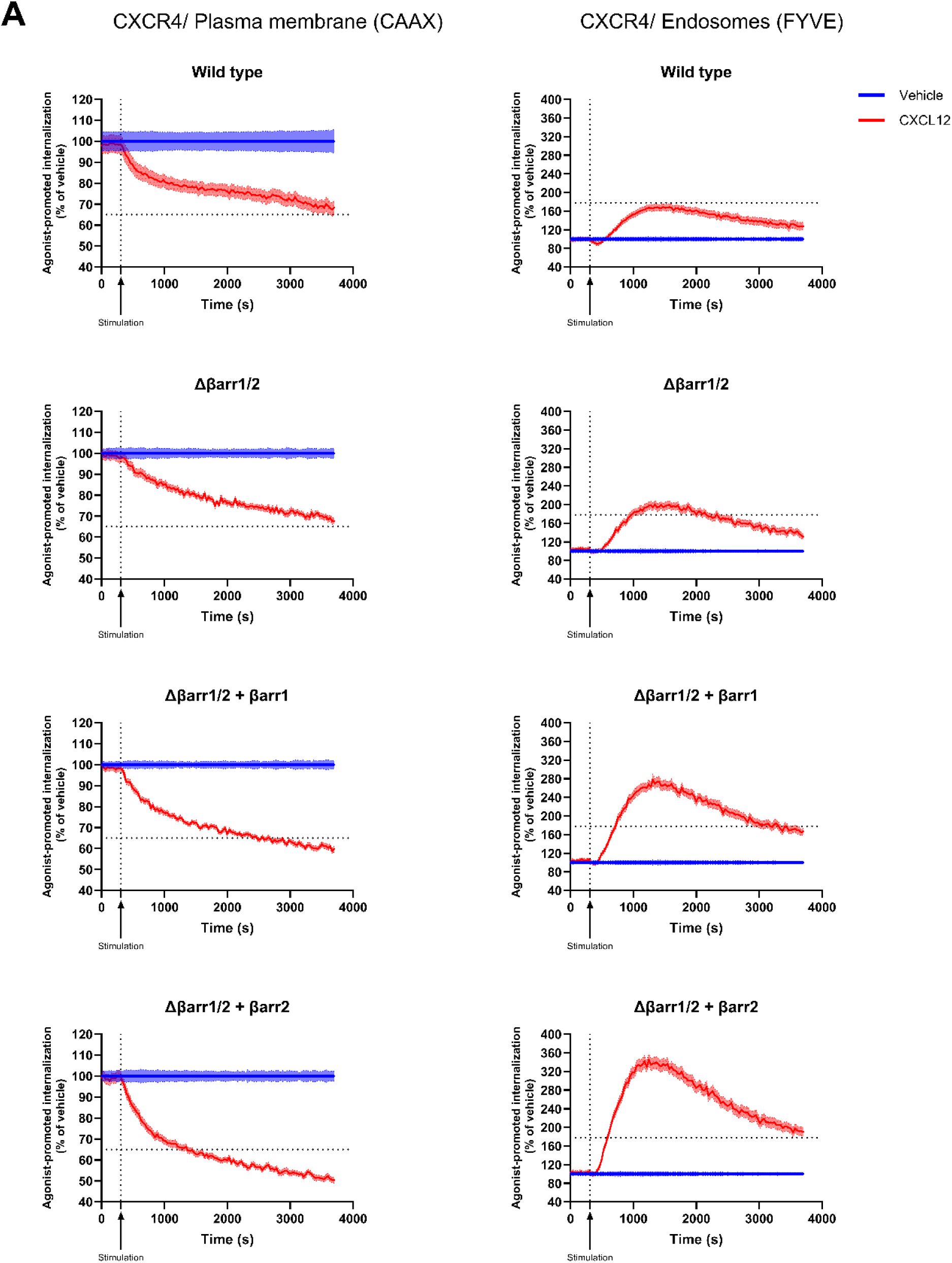
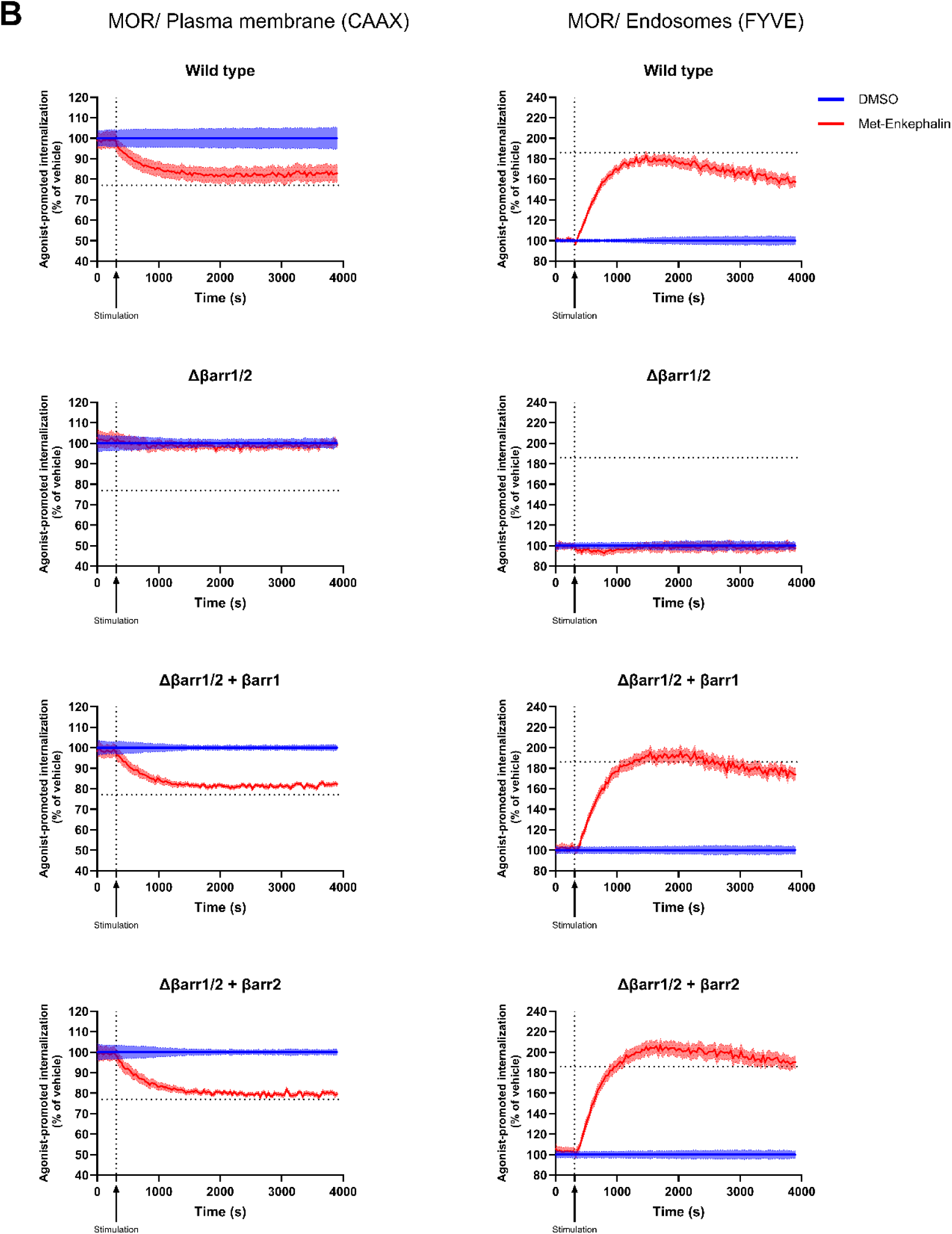
Intracellular trafficking of CXCR4 and MOR in HEK293 cells in the presence or absence of β-arrestins. (**A)** Ligand-induced internalization and early endosome association of CXCR4 were assessed in wild-type HEK293 cells or in cells depleted of β-arrestin isoforms (Δβarr1/2), transfected with plasmids encoding the indicated β-arrestins (βarr1 or βarr2). Cells were co-transfected with plasmids encoding CXCR4-RLucII and either rGFP-CAAX (left panels, CAAX) or rGFP-FYVE (right panels, FYVE). BRET signals were recorded in kinetic mode for 60 min, with ligand stimulation (100 nM CXCL12) applied at 5 min, as indicated by the arrow. Signals were normalized to the vehicle control, set to 100%. (B) Ligand-induced internalization and early endosome association of MOR were assessed in wild-type HEK293 cells or Δβarr1/2 cells transfected with plasmids encoding the indicated β-arrestins (βarr1 or βarr2). Cells were co-transfected with plasmids encoding MOR-RLucII and either rGFP-CAAX (left panels, CAAX) or rGFP-FYVE (right panels, FYVE). BRET signals were recorded for 60 min, with ligand stimulation (5 µM Met-enkephalin) applied at 5 min, as indicated by the arrow. Signals were normalized to the vehicle control, set to 100%. For both panels, the maximal response observed in wild-type cells is indicated by a horizontal line and was used as a reference for β-arrestin–depleted cells (Δβarr1/2) and β-arrestin rescue conditions (Δβarr1/2 + βarr), allowing direct comparison across conditions. All data are presented as mean ± SEM of triplicates from three independent experiments.

**Supplementary Figure 3.**
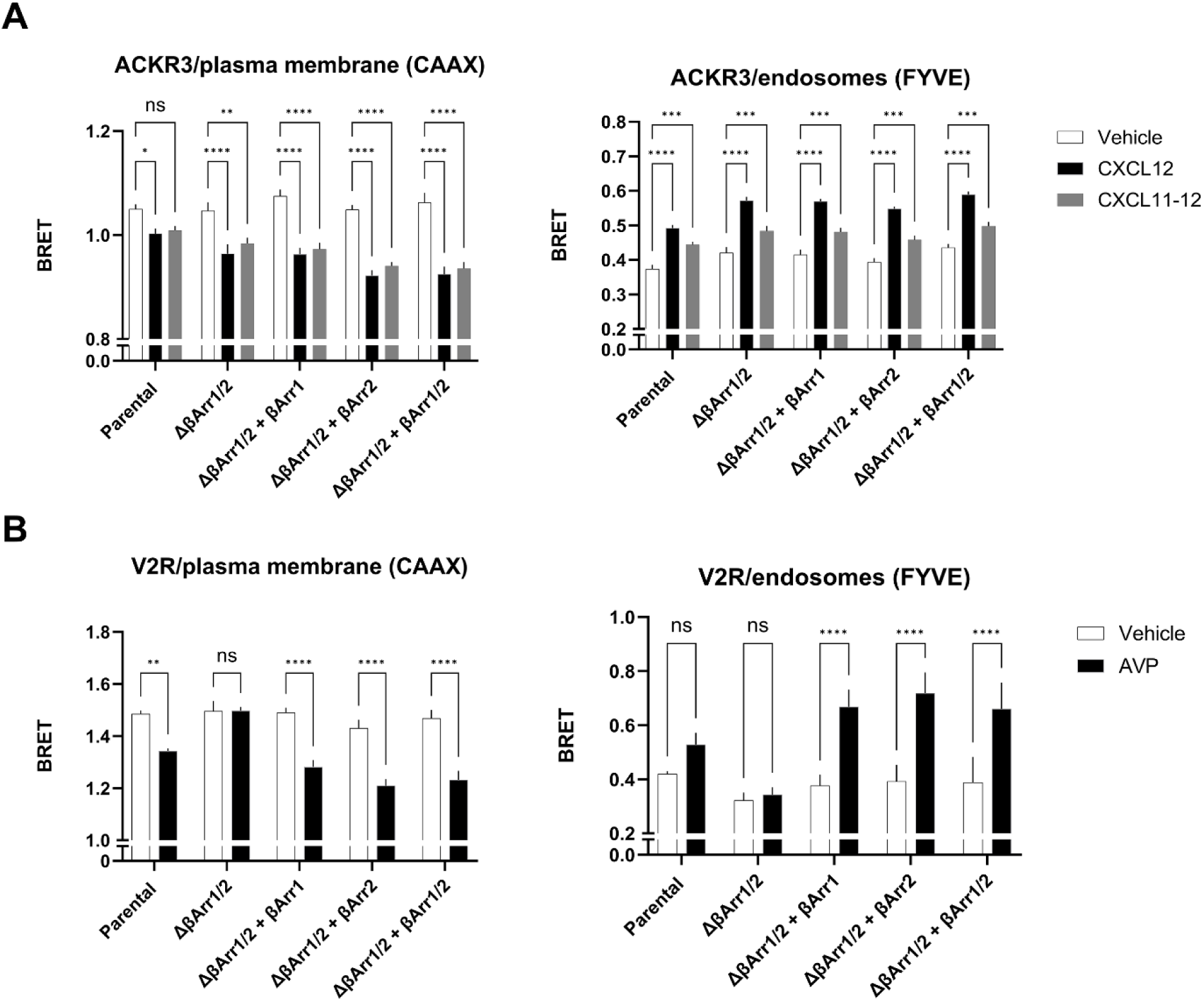
Intracellular trafficking of ACKR3 and V2R in HeLa cells in the presence and absence of β-arrestins. **(A)** Ligand induced internalization and early endosome association of ACKR3 were assessed in HeLa cells (parental) or depleted for β-arrestin isoforms (Δβarr1/2) transfected with plasmids encoding the indicated β-arrestins (βarr1, βarr2 or both) and ACKR3-RLucII and rGFP-CAAX (left panel, CAAX) and ACKR3-RLucII and rGFP-FYVE (right panel, FYVE). Cells were incubated for 25 min in the absence (white bars), the presence of 50 nM CXCL12 (black bars) or 50 nM CXCL11-12 (grey bars). Data are presented as mean ± SEM of triplicates from nine independent experiments. Statistical analysis was performed using two-way ANOVA followed by Dunnett’s multiple comparisons test (ns: not significant, *p<0.05, **p<0.01, ***p<0.001, ****p<0.0001). (B) Ligand induced internalization and association with early endosomes of V2R were assessed in HeLa cells (parental) or depleted for β-arrestins (Δβarr1/2) transfected with plasmids encoding V2R-RLucII and the subcellular compartment markers rGFP-CAAX (left panel, CAAX) and rGFP-FYVE (right panel, FYVE). Cells were incubated for 25 min in the absence (white bars) or presence of 100 nM AVP (black bars). Data are presented as mean ± SEM of triplicates from five independent experiments. Statistical analysis was performed using two-way ANOVA followed by Šídák’s multiple comparisons test (ns: not significant, **p<0.01, ****p<0.0001).

**Supplementary Figure 4.**
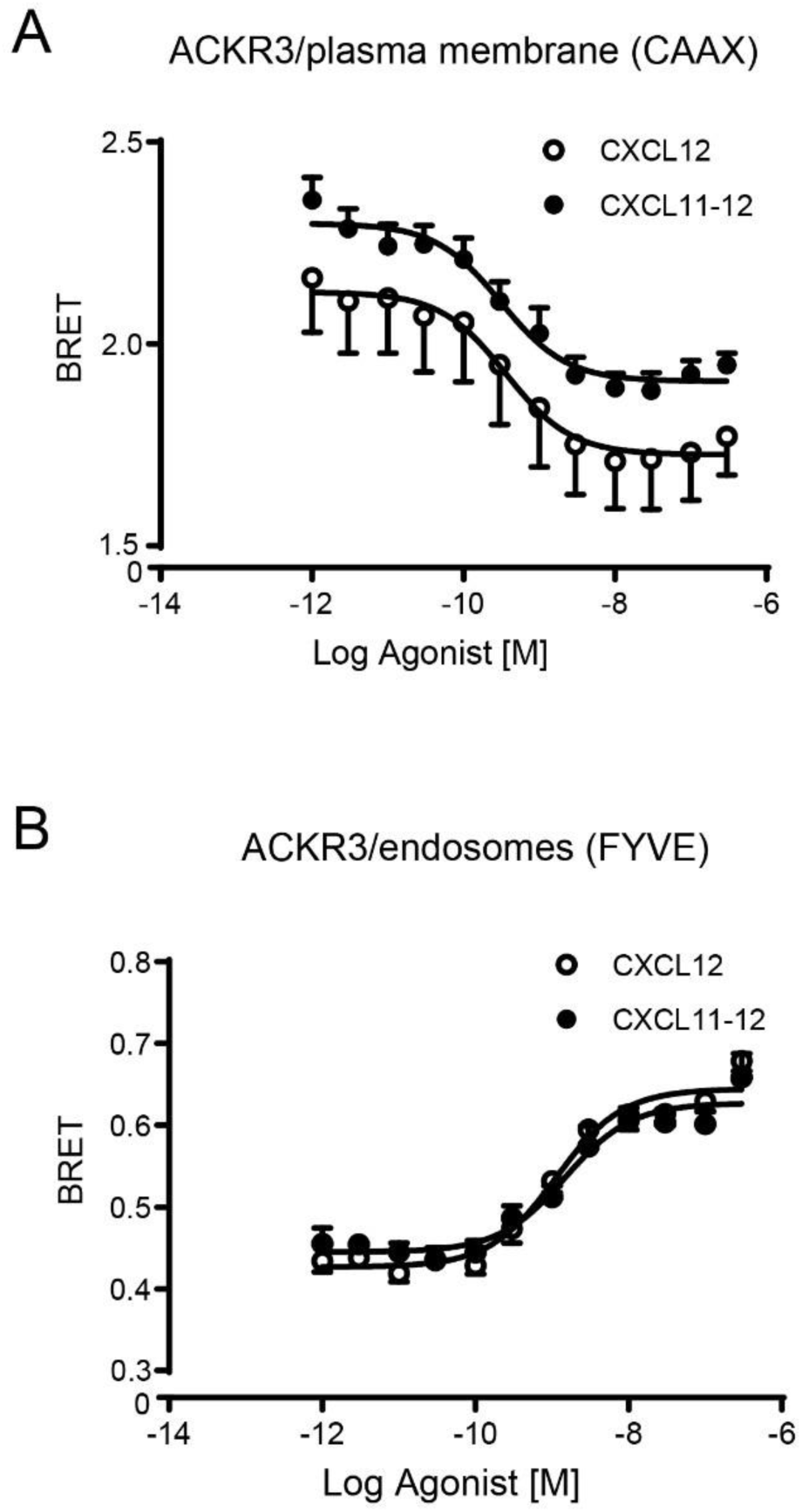
Agonists dependent internalization of ACKR3 in HEK293 cells. HEK293 cells transfected with plasmids encoding ACKR3-RlucII together with either rGFP-CAAX (**A**) or rGFP-FYVE (**B**) were stimulated with the indicated concentrations of CXCL12 or the ACKR3 selective ligand CXCL11-12 for 10 min while BRET signals were recorded. **(A)** Ligand induced dissociation of ACKR3 from the plasma membrane marker rGFP-CAAX in concentration dependent manner, with EC_50_ = 0.36 ± 0.45 and 0.31 ± 0.18 nM for CXCL12 and CXCL11-12 respectively. (B) Ligand induced association of ACKR3 with rGFP-FYVE^+^ early endosomes in concentration dependent manner, with EC_50_ = 1.12 ± 0.10 and 1.48 ± 0.14 nM for CXCL12 and CXCL11-12 respectively. All data are presented as mean ± SEM of triplicates from three independent experiments.

**Supplementary Figure 5.**
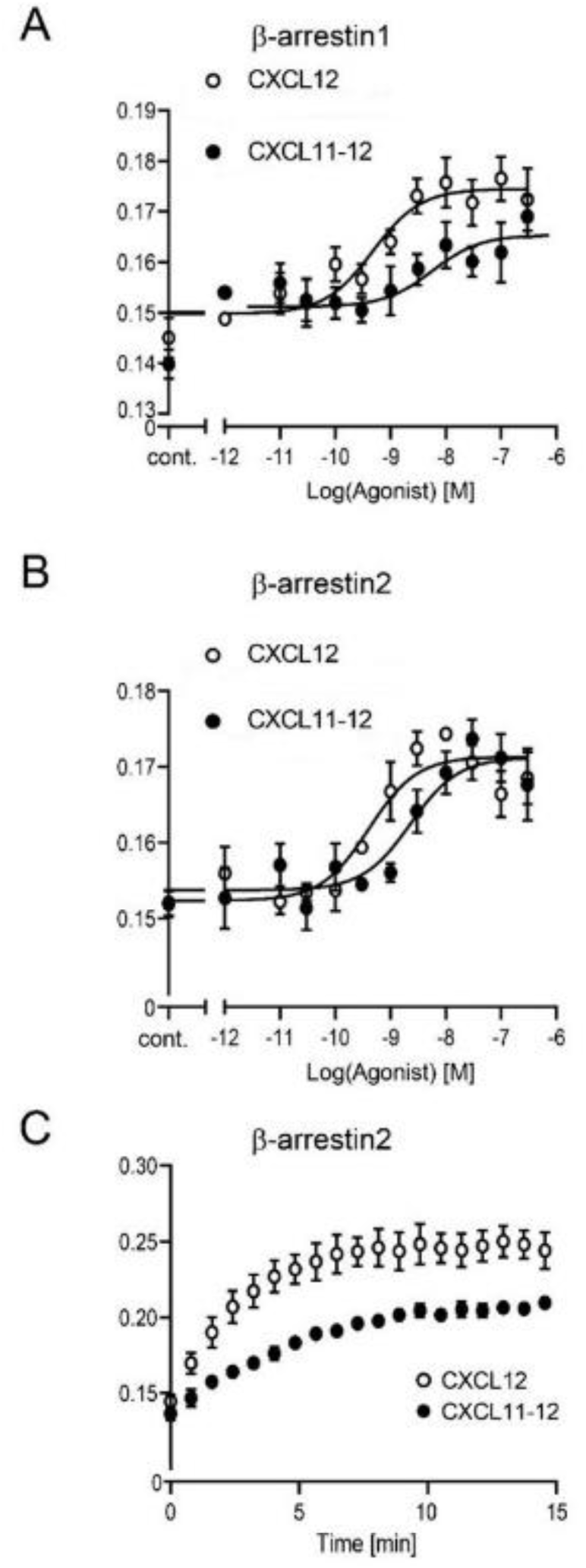
Agonist-dependent recruitment of β-arrestins to the plasma membrane in HeLa cells. HeLa cells were transfected with plasmids encoding ACKR3 together with rGFP-CAAX and β-arrestin1-RlucII (**A**) or β-arrestin-2 RlucII (**B**). Cells were stimulated for 10 min with the indicated concentrations of CXCL12 or the ACKR3 selective ligand CXCL11-12 while BRET signals were recorded. Ligand induced both β-arrestins recruitment to ACKR3 with EC_50_ for β-arrestin1 (CXL12 = 0.47 ± 0.08 nM; CXCL11-12 = 2.23 ± 0.89 nM) and for β-arrestin2 (CXL12 = 0.41 ± 0.05 nM; CXCL11-12 = 2.3 ± 0.42 nM). Data are presented as mean ± SEM of duplicates from six independent experiments. (C) Kinetics of ligand-induced β-arrestin2 recruitment to the plasma membrane in HeLa cells transfected with plasmids encoding ACKR3, rGFP-CAAX and β-arrestin2-RlucII. Cells were stimulated for 15 min with 50 nM of CXCL12 or CXCL11-12. BRET signals were recorded at 30 second intervals. Data are presented as mean ± SEM of triplicates from three independent experiments.

**Supplementary Figure 6.**
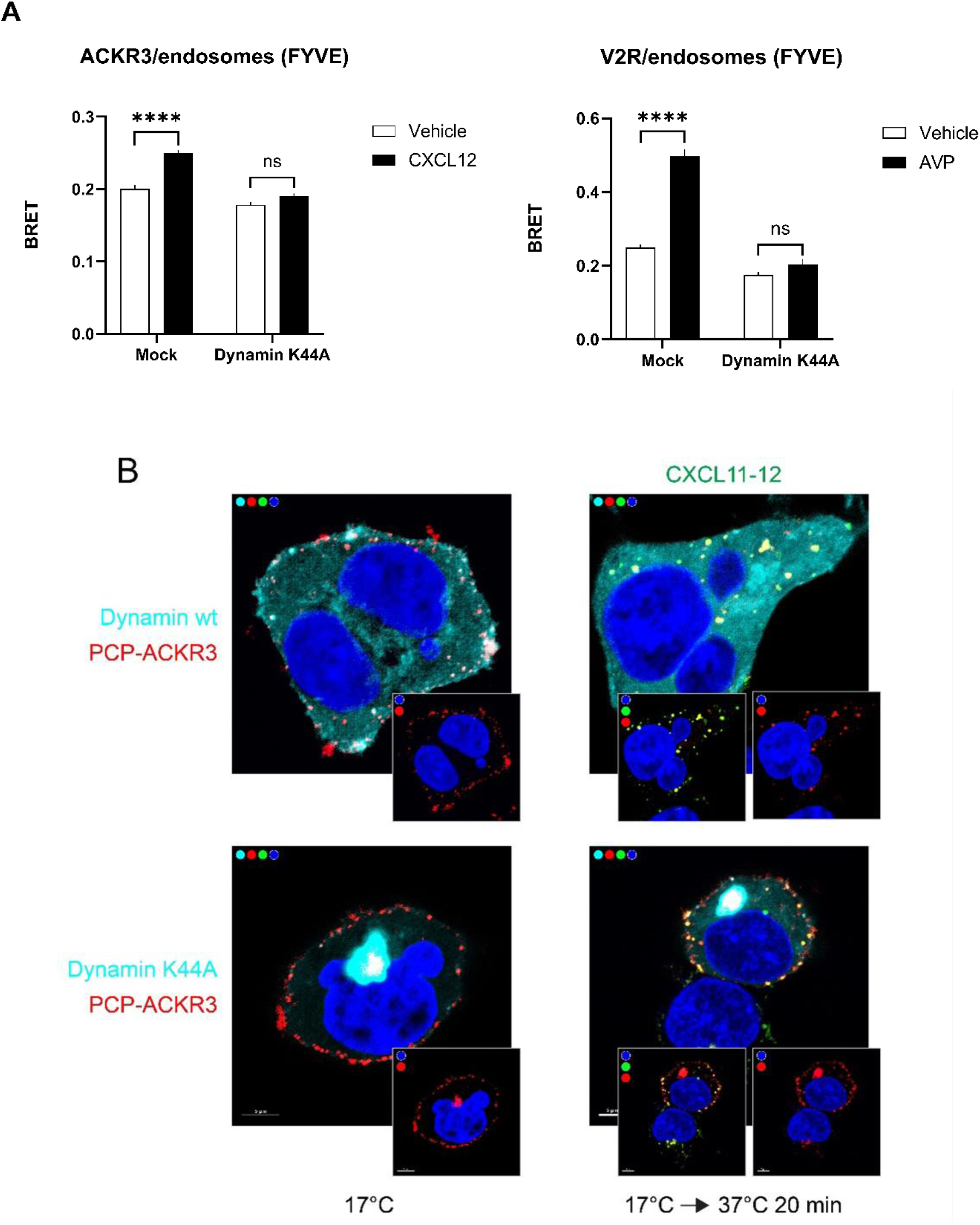
ACKR3 internalization and chemokine scavenging are dynamin dependent. **(A)** HEK293 cells were transfected with plasmids encoding ACKR3-RLucII (left panel) or V2R-RLucII (right panel) along with rGFP-FYVE and, where indicated, together with dominant negative dynamin (Dynamin K44A) or not (Mock). Cells were stimulated with CXCL12 (100 nM, left panel) or arginine-vasopressin (AVP 100nM, right panel) or with medium alone (Vehicle). Data are presented as mean ± SEM of duplicates from three (left panel) and four (right panel) independent experiments. Statistical analysis was performed using unpaired two-tailed T-test; **** p < 0.0001, ns: not significant. (B) HEK293 cells were transiently transfected with PCP-ACKR3 and labeled (red) (at 17°C, as in Figure 1), wild type dynamin (Dynamin wt, cyan, upper panels) or dominant negative dynamin (Dynamin K44A, cyan, lower panels). Cells were shifted to 37°C for 20 min in the presence of 50 nM CXCL11-12Atto565 (green). For clarity in the smaller frames (lower parts) the cyan or chemokine (right) channels were turned off showing localization of PCP-ACKR3 (red) and CXCL-11-12Atto565 (green) (colocalization yellow). Scale bar 5 µm.

**Supplementary Figure 7:**
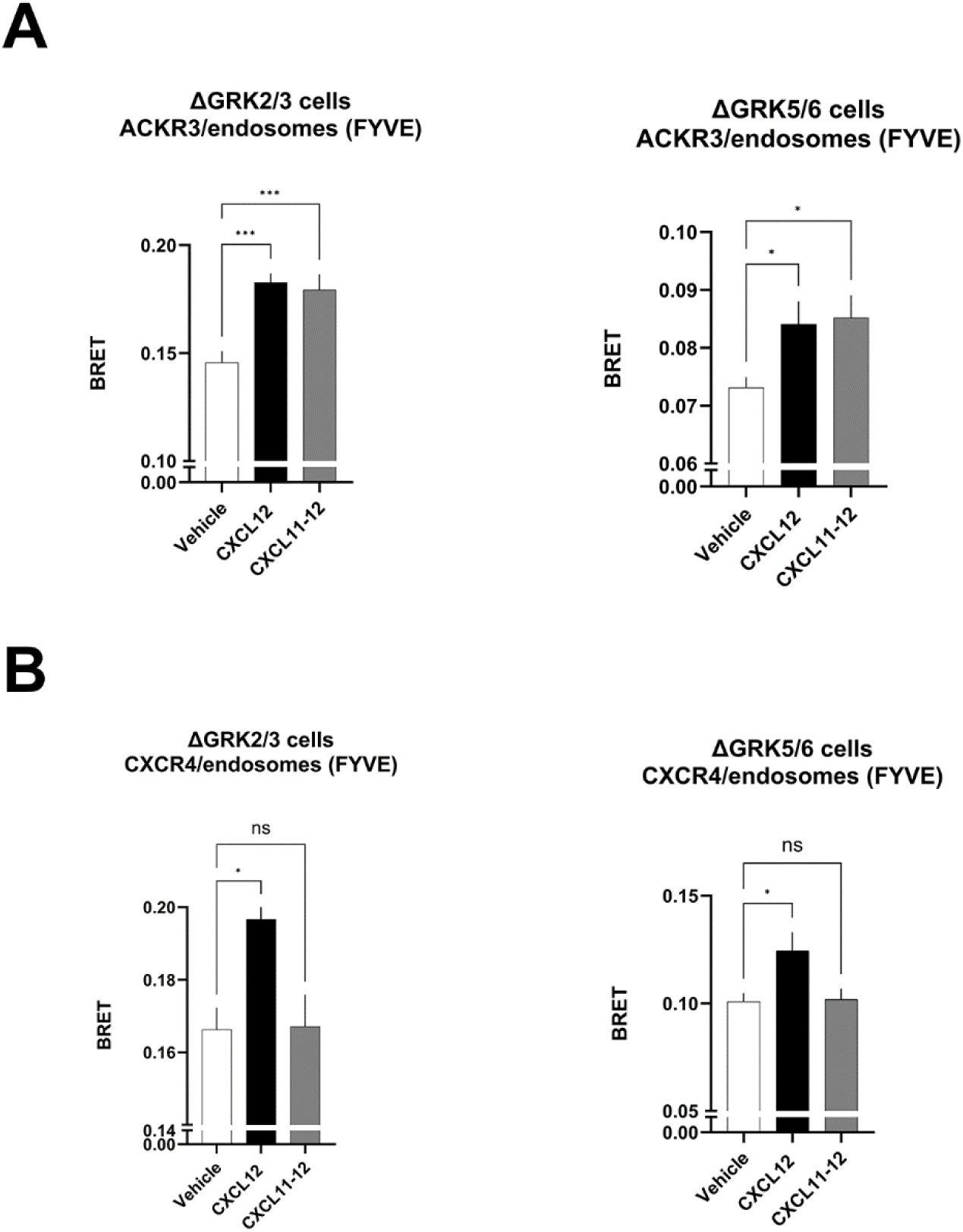
Ligand-dependent ACKR3 and CXCR4 trafficking in the presence or absence of endogenous GRK2/3 and GRK5/6. **(A)** Ligand induced early endosome association of ACKR3 was assessed in HEK293 cells depleted for GRK2 and GRK3 (ΔGRK2/3 cells, left panel) or depleted for GRK5 and GRK6 (ΔGRK5/6 cells, right panel) transfected with plasmids encoding ACKR3-RLucII and rGFP-FYVE (FYVE). Cells were incubated for 25 min in the absence (white bars), the presence of 100 nM CXCL12 (black bars) or 100 nM CXCL11-12 (grey bars). (B) Ligand induced early endosome association of CXCR4 was assessed in HEK293 cells depleted for GRK2 and GRK3 (ΔGRK2/3 cells, left panel) or depleted for GRK5 and GRK6 (ΔGRK5/6 cells, right panel) transfected with plasmids encoding CXCR4-RLucII and rGFP-FYVE (FYVE). Cells were incubated for 25 min in the absence (white bars), the presence of 100 nM CXCL12 (black bars) or 100 nM CXCL11-12 (grey bars). All data are presented as mean ± SEM of triplicates from four independent experiments. Statistical analysis was performed using one-way ANOVA followed by Dunnett’s multiple comparisons test (ns: not significant, *p<0.05, ***p<0.001).

**Supplementary Figure 8.**
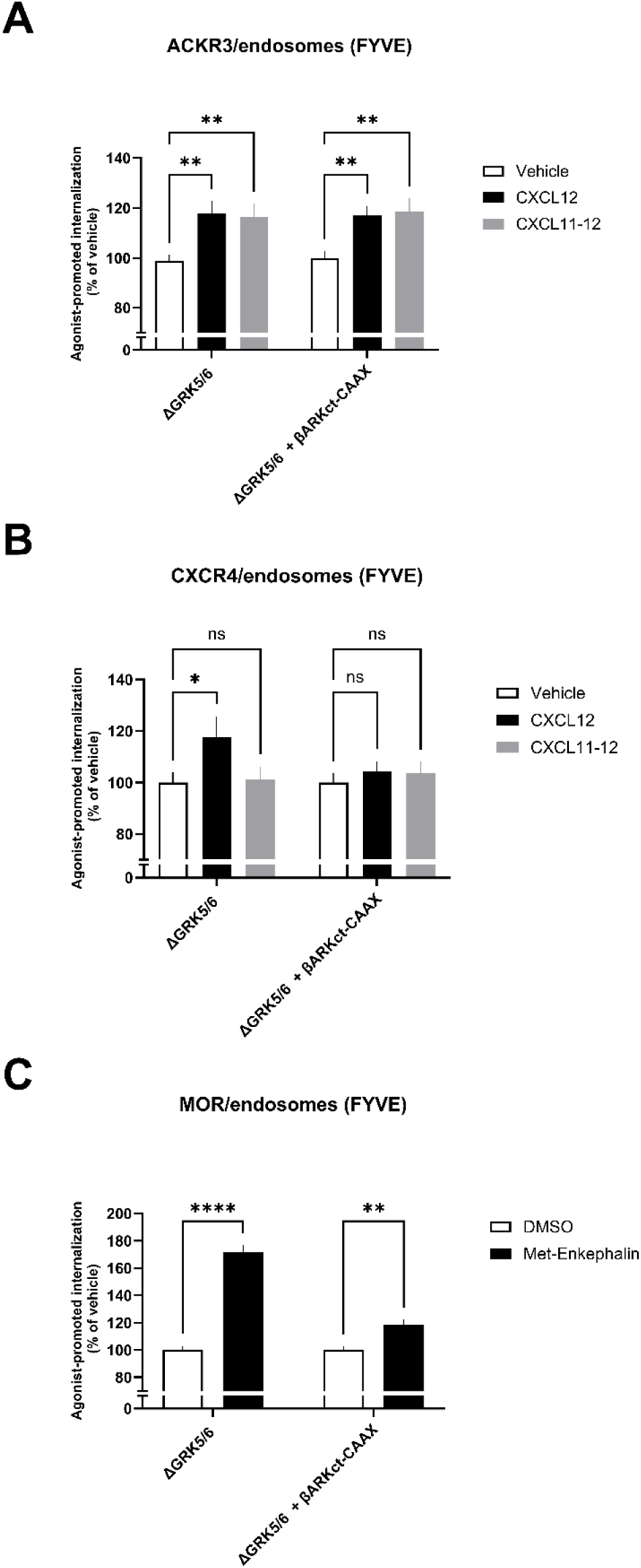
Effect of Gβγ blocker (βARKct-CAAX) in GRK2 promoted endocytosis of ACKR3, CXCR4 and MOR in ΔGRK5/6 cells. **(A)** Ligand-induced early endosome association of ACKR3 was assessed in HEK293 cells depleted for GRK5 and GRK6 (ΔGRK5/6), transfected or not with a plasmid encoding βARKct-CAAX. Cells were co-transfected with plasmids encoding ACKR3-RLucII and rGFP-FYVE. Cells were incubated for 25 min in the absence of ligand (white bars), or in the presence of 100 nM CXCL12 (black bars) or 100 nM CXCL11-12 (grey bars). Statistical analysis was performed using two-way ANOVA followed by Dunnett’s multiple comparison test (**p < 0.01). (B) Ligand-induced early endosome association of CXCR4 was assessed in HEK293 cells depleted for GRK5 and GRK6 (ΔGRK5/6), transfected or not with a plasmid encoding βARKct-CAAX. Cells were co-transfected with plasmids encoding CXCR4-RLucII and rGFP-FYVE. Cells were incubated for 25 min in the absence of ligand (white bars), or in the presence of 100 nM CXCL12 (black bars) or 100 nM CXCL11-12 (grey bars). Statistical analysis was performed using two-way ANOVA followed by Dunnett’s multiple comparison test (ns, not significant; *p < 0.05). (C) Ligand-induced early endosome association of MOR was assessed in HEK293 cells depleted for GRK5 and GRK6 (ΔGRK5/6), transfected or not with a plasmid encoding βARKct-CAAX. Cells were co-transfected with plasmids encoding MOR-RLucII and rGFP-FYVE. Cells were incubated for 25 min in the absence of ligand (white bars) or in the presence of 5 µM Met-enkephalin (black bars). Statistical analysis was performed using two-way ANOVA followed by Šídák’s multiple comparison test (**p < 0.01; ****p < 0.0001). All data are presented as mean ± SEM of triplicates from four independent experiments.

**Supplementary Figure 9.**
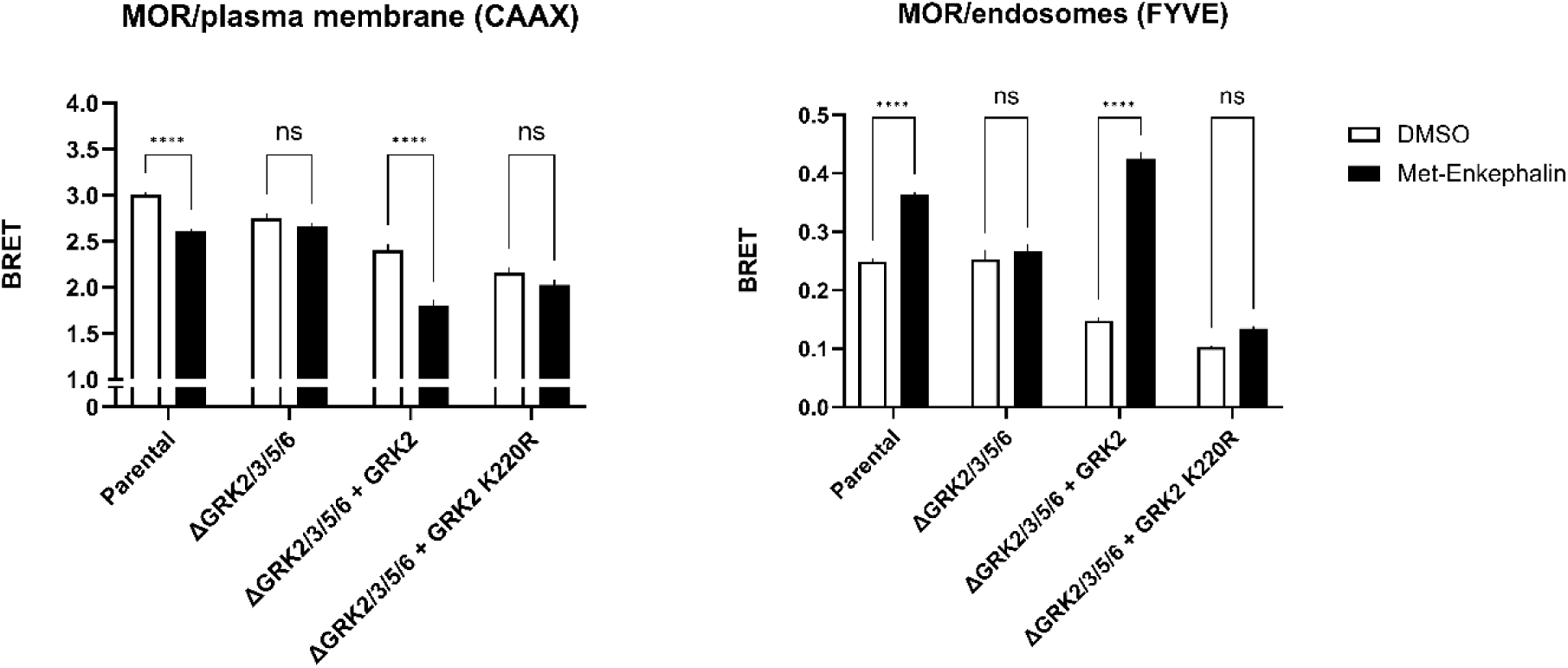
Ligand-dependent MOR trafficking in the presence or absence of kinase-dead GRK2 (K220R). Ligand-induced internalization and early endosome association of MOR were assessed in parental HEK293 cells or in cells depleted of GRK2, GRK3, GRK5, and GRK6 (ΔGRK2/3/5/6), transfected or not with a plasmid encoding GRK2 or GRK2 K220R (kinase-dead mutant). Cells were co-transfected with plasmids encoding MOR-RLucII and either rGFP-CAAX (left panel, CAAX) or rGFP-FYVE (right panel, FYVE). Cells were incubated for 25 min in the absence of ligand (white bars) or in the presence of 5 µM Met-enkephalin (black bars). Data are presented as mean ± SEM of triplicates from three independent experiments. Statistical analysis was performed using two-way ANOVA followed by Sidak’s multiple comparison test (ns, not significant; ****p < 0.0001).

**Supplementary Figure 10.**
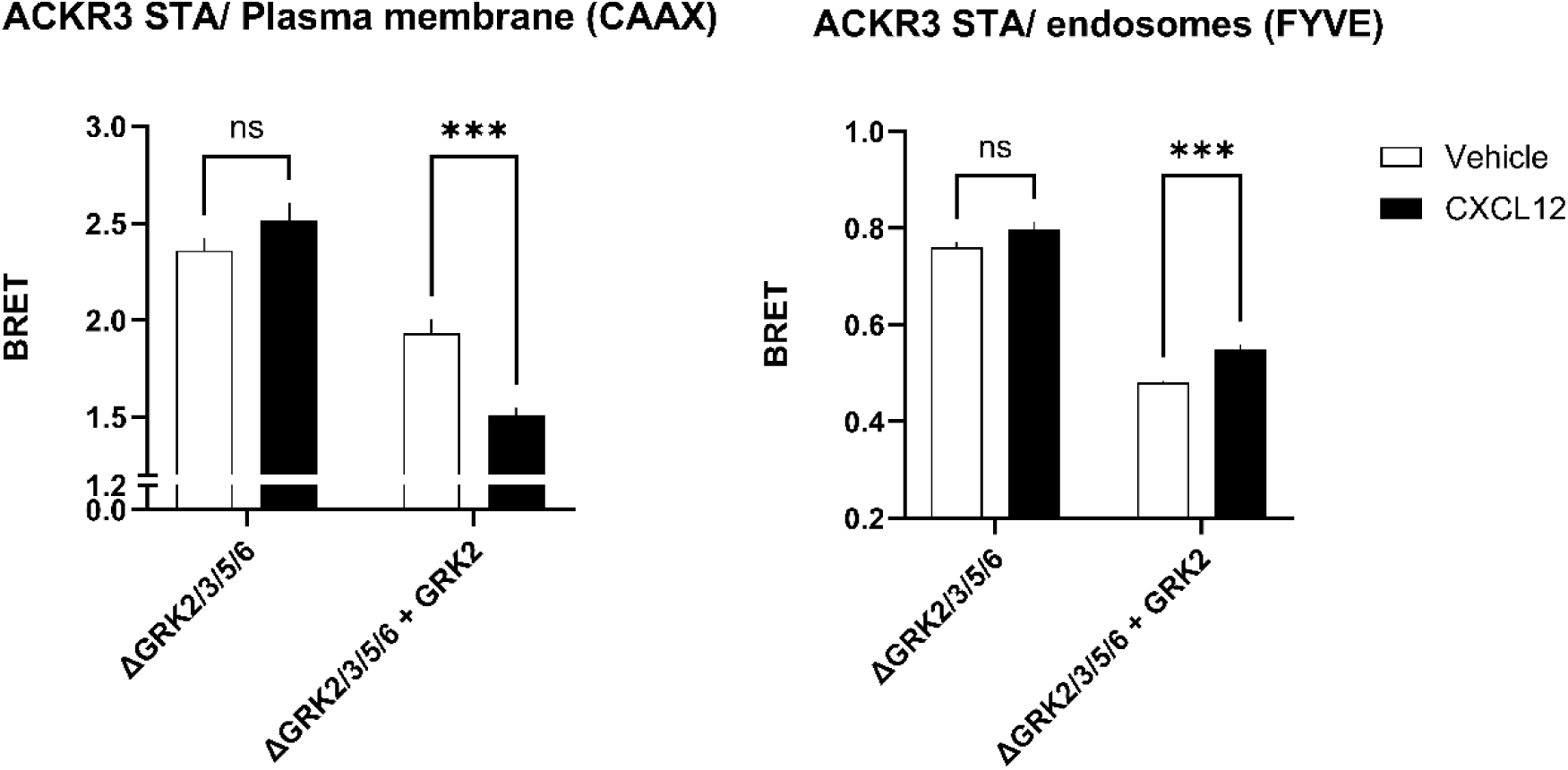
Ligand-induced ACKR3 STA endocytosis in the presence or absence of GRK2. Ligand induced internalization of ACKR3 phosphodeficient mutant (ST/A) in HEK293 cells depleted for GRK2, GRK3, GRK5, GRK6 (ΔGRK2/3/5/6) transfected or not with a plasmid encoding GRK2. Cells were co-transfected with plasmids encoding ACKR3-ST/A-RLucII and either rGFP-CAAX (left panel, CAAX) or rGFP-FYVE (right panel, FYVE). Cells were incubated for 25 min in the absence (white bars) or the presence of 100 nM CXCL12 (black bars). Data are presented as mean ± SEM of triplicates from three independent experiments. Statistical analysis was performed using two-way ANOVA followed by Sidak’s multiple comparison test (ns: not significant, ***p<0.001).

